# Tau modulates mRNA transcription, alternative polyadenylation profiles of hnRNPs, chromatin remodeling and spliceosome complexes

**DOI:** 10.1101/2021.07.16.452616

**Authors:** Mauro Montalbano, Elizabeth Jaworski, Stephanie Garcia, Anna Ellsworth, Salome McAllen, Andrew Routh, Rakez Kayed

**Affiliations:** Mitchell Center for Neurodegenerative Diseases, University of Texas Medical Branch, Galveston, Texas, 77555, USA; Departments of Neurology, Neuroscience and Cell Biology, University of Texas Medical Branch, Galveston, Texas, 77555, USA; Department of Biochemistry and Molecular Biology, University of Texas Medical Branch, Galveston, Texas 77555, USA; Sealy Center for Structural Biology and Molecular Biophysics, University of Texas Medical Branch, Galveston, TX, USA

**Keywords:** Tau, Transcriptomic, Alternative Polyadenilation, Nuclear Dysfunction, Neurodegeneration

## Abstract

Tau protein is a known contributor in several neurodegenerative diseases, including Alzheimer ‘s disease (AD) and frontotemporal dementia (FTD). It is well-established that tau forms pathological aggregates and fibrils in these diseases. Tau has been observed within the nuclei of neurons, but there is a gap in understanding regarding the mechanism by which tau modulates transcription. We are interested in the P301L mutation of tau, which has been associated with FTD and increased tau aggregation. Our study utilized tau-inducible HEK (iHEK) cells to reveal that WT and P301L tau distinctively alter the transcription and alternative polyadenylation (APA) profiles of numerous nuclear precursors mRNAs, which then translate to form proteins involved in chromatin remodeling and splicing. We isolated total mRNA before and after over-expressing tau and then performed Poly(A)-ClickSeq (PAC-Seq) to characterize mRNA expression and APA profiles. We characterized changes in Gene Ontology (GO) pathways using EnrichR and Gene Set Enrichment Analysis (GSEA). We observed that P301L tau up-regulates genes associated with reactive oxygen species responsiveness as well as genes involved in dendrite, microtubule, and nuclear body/speckle formation. The number of genes regulated by WT tau is greater than the mutant form, which indicates that the P301L mutation causes loss-of-function at the transcriptional level. WT tau up-regulates genes contributing to cytoskeleton-dependent intracellular transport, microglial activation, microtubule and nuclear chromatin organization, formation of nuclear bodies and speckles. Interestingly, both WT and P301L tau commonly down-regulate genes responsible for ubiquitin-proteosome system. In addition, WT tau significantly down-regulates several genes implicated in chromatin remodeling and nucleosome organization. Although there are limitations inherent to the model systems used, this study will improve understanding regarding the nuclear impact of tau at the transcriptional and post-transcriptional level. This study also illustrates the potential impact of P301L tau on the human brain genome during early phases of pathogenesis.

**Author summary:** While tau biology has been extensively studied and closely linked to several neurodegenerative diseases, our current understanding of tau’s functions in the nucleus is limited. Given the role of tau in disease progression and pathogenesis, elucidating the function of tau activity in transcription and its nuclear accumulation may reveal novel therapeutic targets; therefore, helping identify new upstream pathways that have yet to be investigated. In this study, we used tau-inducible cell lines to uncover new molecular mechanisms by which tau functions in the nucleus. This study systematically investigates the changes in transcriptomic and alternative polyadenylation profiles modulated by WT and mutant P301L tau protein. In this manuscript, we report following new findings (**i**) tau modulates gene expression of transcripts associated with chromatin remodeling and splicing complexes; (**ii**) WT and mutant P301L tau regulate, differentially, transcription and alternative polyadenylation (APA) profiles; and (**iii**) P301L mutation affects the transcription mediated by tau protein. The potential role of tau in mediating transcription and alternative polyadenylation processes is not well studied, representing a novelty in the field. Therefore, this research establishes a new direction for investigating tau nuclear function in both human and mouse brains.

## Introduction

Tau is a neuronal protein found both inside and outside of the nucleus that contributes to the pathology of neurodegenerative diseases such as frontotemporal dementia (FTD) and Alzheimer’s disease (AD)^1^. It is primarily described as a microtubule-associated protein^2^. Nuclear tau has been found to ‘protect’ DNA^1–3^ during reactive oxygen species (ROS)-induced heat stress. However, nuclear and cytosolic tau interact with RNA to form droplets^4^ and aggregates^5^. Tau has also been observed altering nuclear structure^6,7^ in the human nuclei of neuroblastoma^8,9^ and in HEK-293 cells. More specifically, phosphorylation of nuclear tau negatively regulates its nuclear function in pluripotent neuronal cells and neuroblastoma cells^10^. Previous studies have revealed that nuclear tau plays a role in the DNA damage response (DDR) through deadenylation, which triggers major mRNA decay pathways^11,12^. Most recently, we found that oligomeric assemblies of tau containing RNA-binding proteins impair chromatin remodeling and nuclear lamina formation through associations with histones and chromatin components in the nuclear compartment^13^.

Despite the well-established importance of tau in the cytoskeleton of neurons^14^, there is growing evidence that tau is notably involved in nucleolar transcription and cellular stress responses^15,16^. Recently, it was shown that mutations and/or the phosphorylation of tau results in the deformation of the neuronal nuclear membrane and can disrupt nucleocytoplasmic transport^17^ in FTD^7,18^ and AD^19,20^. Related studies analyzed the direct impact in transcriptional activity due to tau and found that nuclear tau regulates the expression of VGluT1, a gene that controls glutamatergic synaptic transmission, and that tau displacement from microtubules (MTs) increases nuclear accumulation of tau^21^. Furthermore, tau modifies histone acetylation and was shown to have a broad epigenomic impact in the aging and pathology of AD human brains^22^. It has also been observed that tau interacts with neuronal pericentromeric DNA regions, particularly in association with HP1 and H3K9me3^23^, this observation spots tau protein as potential chromatin remodeling factor. Lastly, tau exhibits binding interactions with genic and intergenic DNA sequences of primary cultured neurons, especially in positions ±5000 bp away from the start site of transcription ^24^.

In eukaryotic cells, the maturation of 3’ ends in mRNA involves endonucleolytic cleavage of the nascent RNA followed by the synthesis of a poly(A) tail on the 3’ terminus of the cleaved product by a poly(A) polymerase (PAP)^25^. This reaction is called polyadenylation and is fundamentally linked to transcription termination. The sequences for the mRNA precursors and the proteins required for polyadenylation are well understood. It has been clearly elucidated that a single gene can give rise to many possible transcripts, each with different polyadenylation sites (poly(A)-sites, or PASs), and that differential usage of these sites can lead to the formation of mRNA isoforms. This phenomenon is called alternative polyadenylation (APA)^26^ and is a common event in eukaryotic cells. In fact, researchers have determined that 50% of mammalian mRNA-encoding genes express APA isoforms^27,28^. Considering this information, we used tau inducible HEK (iHEK) cell lines to obtain and analyze transcriptomic and APA profiles in the presence of WT and P301L tau. To characterize transcriptional and post-transcriptional profiles modified by WT and P301L, we utilized Poly(A)-ClickSeq (PAC-Seq) to measure changes in the expression of the host mRNA transcript whilst simultaneously characterizing changes in the PAS usage or creation of mRNA isoforms. In addition, we employed Gene Set Enrichment Analysis (GSEA) and Gene Ontology (GO) to study the main gene domains modulated by tau.

## Materials and Methods

### Cell Culture and Tau Expression

In this study we used two different versions of tau inducible HEK (iHEK) cells: iHEK overexpressing WT tau and iHEK overexpressing mutated P301L tau. They were maintained in Dulbecco’s modified eagle medium (DMEM) supplemented with 10% fetal bovine serum (FBS) at 37 oC in 5% CO_2_. To induce WT and mutant tau overexpression, iHEK cells were treated with 1µg/mL of Tetracycline (Tet) for 24 hours in FBS-depleted DMEM (Gibco™ LS11965118, Fisher Scientific). iHEK cells not treated with Tet were named control (Ctr). After 24 hours, two washes with medium were done to remove excess Tet. Immediately after the washes, the cells were stained and collected. Detachment of cells was completed with Trypsin (Gibco™ Trypsin-EDTA, 0.25% Phenol red, LS25200114 Fisher Scientific), and the cells warmed for 3 minutes in the incubator following the addition of Trypsin. The cells were then centrifuged at 1000 rpm for 5 minutes. Lastly, cell pellets were harvested and used for protein fractionation, and mRNA extraction.

### RNA Extraction

Total mRNA was collected by using TRIzol extraction reagent according to established protocol^29^. RNA samples for Real Time Analysis (RT-PCR) were quantified using a Nanodrop Spectrophotometer (Nanodrop Technologies), followed by analysis on an RNA Nano chip using the Agilent 2100 Bioanalyzer (Agilent Technologies). Only samples with high quality total RNA were used (RIN: 7.5-10.0) for the study. Synthesis of cDNA was performed with either 0.5µg or 1µg of total RNA in a 20µl reaction using the reagents available within the Taqman Reverse Transcription Reagents Kit from Life Technologies (#N8080234). Q-PCR amplifications (performed in duplicate or triplicate) were done using 1µl of cDNA in a total volume of 20µl using the iTaq Universal SYBR Green Supermix (Bio-Rad #1725125). The final concentrations of the primers were 300nM. Relative RT-QPCR assays are performed with either 18S RNA gene as a normalizer. Absolute RNA quantification analysis was performed using known amounts of a synthetic transcript created from the gene of interest.

### Library Preparation Protocol

Protocols for Poly(A)-ClickSeq (PAC-Seq) have been described in detail by Jaworski et al. 2018 ^30,31^. Approximately 1µg of total cellular RNA per sample was used as a template in reverse-transcription reactions supplemented with 40uM Azido-VTPs and primed using an oligo-dT primer containing a partial Illumina i7 indexing adaptor. Azido-terminated cDNA fragments were ‘click-ligated’ to hexynyl-functionalized click-adaptors containing the Illumina i5 universal sequencing adaptor. Single-stranded cDNA libraries were indexed in a final PCR reaction for 15-18 PCR cycles. Final libraries were size extracted by gel-electrophoresis and submitted for sequencing using an Illumina NextSeq550 to prepare 1×150 SE reads. RNAseq datasets is uploaded to NCBI SRA, reference number: <u>PRJNA744518</u>.

### Poly(A)-ClickSeq

PAC-Seq data were analyzed using the Differential Poly-A Clustering (*DPAC*) program, which ran with default settings as previously described^32^. *DPAC* trims and quality-filters raw FASTQ data and therefore requires each read to have at least 25 ‘As’ at the 3’ end of the read. These reads are then trimmed using *cutadapt*. Trimmed reads are mapped to the reference human genome (hg19) using *HISAT2*^33^. The 3’end of mapped reads are thus used to annotate poly(A)-sites and annotated based upon overlaps with gene annotations obtained from UCSC genome browser. Gene counts were extracted and *DESeq2* was used to calculated changes in gene expression as well as relative changes in expression in individual poly(A)-sites found within single genes. Differential gene expression was assigned when a gene had a fold change greater than +/- 1.5-fold with a p-adj value less than 0.1. Alternative polyadenylation is assigned when a single gene has two or more clustered poly(A)-sites wherein at least one of these sites has a differential usage greater than a +/- 1.5-fold, a p-adj value less than 0.1, and a change of the relative usage of a poly(A)-cluster within the gene of greater than 10%.

### Western Blotting and Cell Fractioning

Immunoblot (IB) analyses were performed with iHEK cell fraction samples as previously described^13^. Approximately 10 µg of protein preparations were loaded onto precast NuPAGE 4-12% Bis-Tris gels (NP0335BOX, Invitrogen) for sodium dodecyl sulfate-polyacrylamide gel electrophoresis (SDS-PAGE) analyses. Gels were subsequently transferred onto nitrocellulose membranes and blocked overnight at 4°C with 10% nonfat dry milk. Membranes were then probed for 1 hour at room temperature with Pan-Tau (Tau13, 1:10,000, MMS-520R Covance), (GAPDH, 1:1000, ab9485 Abcam), Histone3 (1:1000, ab201456 Abcam), RCC1 (1:100, Clone E-6 sc-55559 Santa Cruz Tech.), DNAJC2 (1:5000, ab134572 Abcam), Histone 1.2 (1:500, ab4086 Abcam), HMGB1 (1:500, ab18256 Abcam), SMARCA5 (1:10000, #PA5-78253, Invitrogen), SMARCC1 (0.4µg/mL, #PA5-55058, Invitrogen), and β-Actin (1:5000, #A1978, Sigma Aldrich). Antibodies were diluted in 5% nonfat dry milk. Immunoreactivity was detected using a horseradish peroxidase (HRP)-conjugated anti-rabbit immunoglobulin G (IgG, 1:10,000, NA934 GE Healthcare). Tau13 and Tau5 immunoreactivity were detected using an anti-mouse IgG (1:10,000, NA931 GE Healthcare) diluted in 5% milk. ECL Plus (K-12045-D50, GE Healthcare) was used to visualize protein bands. LaminB1/Histone3 and GAPDH were used to normalize and quantify nuclear and cytoplasmic proteins, respectively. The compartment extraction was conducted with Qproteome Cell Compartment Kits (Qiagen, #37502); nuclear, cell membrane, and cytoplasmic proteins were isolated and preserved for IB analysis.

### Immunofluorescence of Fixed Cells and Fluorescence Microscopy

Cells on a 24-well coverslip were fixed with 0.5 ml of 4% PFA/PBS for 15 min. The cells were then washed 3 times in phosphate buffered saline (PBS), for 5 min for each wash. The cells were permeabilized in 0.5ml PBS and 0.2% Triton X-100 in phosphate buffered saline containing 0.5% Tween (PBST) for 5 min. Blocking was done in 0.5 ml of 5% normal goat serum (NGS) in PBST for 1 hour. Primary antibody was diluted in 5% NGS/PBST overnight at 4°C for incubation, and then washed 3 times in PBST, for 10 min each. Secondary antibody diluted in 5% NGS/PBST was incubated for 2 hours at room temperature. All the secondary antibodies were purchased from Thermo Fisher Scientific and used at a 1:800 dilution for staining. After applying secondary antibodies, cells were incubated in DAPI (nuclei staining) diluted 1:10,000 in PBST (5 mg/ml stock solution) for 5 min after the first wash. The cells were then washed 2 times with PBST, and once with PBS (10 min each) prior to mounting coverslips. Coverslips were mounted on glass microscope slides using 8-10 µl of Prolong Gold Antifade mounting media with DAPI (Invitrogen, P36941) per coverslip. Slides were air-dried in fume hood or stored at 4°C until ready to be dried in the fume hood. The primary antibodies used in this study for immunocytochemistry (ICC) are as follows: Histone 1.2 (Abcam ab4086 - 1 µg/ml), Ki-67 (Abcam ab92742 - 1 µg/ml), SMARCC1 (Invitrogen PA5-55058 - 0.25 µg/ml, SMARCA5 (Invitrogen PA5-78253 - 1 µg/ml, MCM2 (Abcam ab108935 - 1/1000), RCC1 (Santa Cruz, INC. sc-55559 - 1:50), and Tau13 (Bio Legend MMS-520R - 1/200). After three washes with PBS, cells were probed with mouse and rabbit-specific fluorescent-labeled secondary antibodies (1:200, Alexa Fluor 488 and 633, Life Technologies). Single frame images were collected using the Keyence BZ-X 710 Microscope. Images for quantification of area and integrated density were taken in nuclear target areas guided by the DAPI fluorescence. We then performed single extraction analysis using BZ-X Analyzer software (Keyence). We used 200 nuclei per target area and used the Nikon 20X objective for imaging and quantification analysis.

### Statistical Analysis

All in-vitro experiments were performed in at least three biological replicates. All data are presented as means ± SD and were analyzed using GraphPad Prism Software 6.0. Statistical analyses included the Student-t Test or one-way ANOVA followed by Tukey’s Multiple Comparisons Test. Column means were compared using one-way ANOVA with treatment as the independent variable. In addition, group means were compared using two-way ANOVA considering factors for each treatment respectively. When ANOVA showed a significant difference, pair-wise comparisons between group means were examined by the Tukey and Dunnett Multiple Comparison Test.

## Results

### WT tau up-regulates genes associated with cytoskeleton organization and nuclear speckles/bodies

Firstly, we evaluated changes in gene expression profiles upon expression of WT and P301L tau in iHEK cells that were induced with tetracycline (Tet). After 24h of Tet induction, we confirmed tau expression in the cytoplasm and nuclei of iHEK cells (Fig S1A). Total cellular RNA from WT and P301L tau (untreated (Control) and treated (+Tet)) study groups was extracted using TRIzol reagent and by following established protocol^7,13^. RNA was sequenced using Poly(A)-ClickSeq (PAC-Seq) to measure changes in gene expression and poly(A)-site usage^31^. A schematic of the experimental design is provided in Fig 1A. Volcano scatterplots from WT and P301L tau iHEK (Fig 1B and 1C, respectively) demonstrate a substantial difference in the number of genes regulated by WT tau and P301L tau. After Tet induction in the WT tau iHEK cell system, we observed up-regulation of 88 genes and down-regulation of 30 genes (gene names listed in Fig 1D). In the P301L tau iHEK cell system, these numbers dropped to 10 up-regulated genes and only 1 down-regulated (gene names listed in Fig 1E).

**Fig 1.**
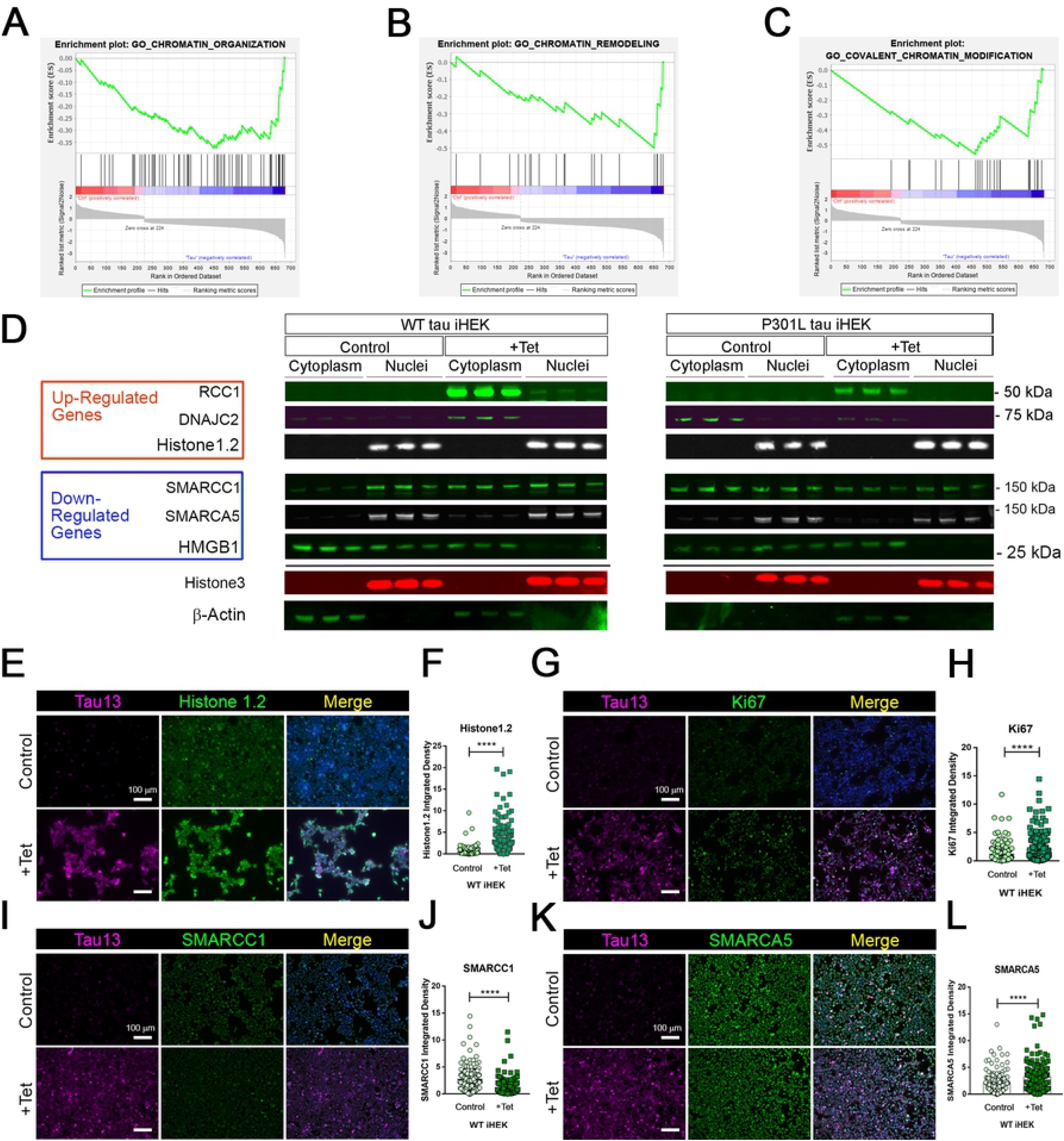
Tau-dependent Gene expression. (**A**) Schematic representation of experimental plan, from Tet induction in WT and P301L Tau IHEK to RNA isolation, sequencing to gene expression analysis. (**B**) Volcano Plot for Down- and Up-regulated gene in WT Tau iHEK. (**C**) Volcano Plot for Down- and Up-regulated gene in P301L Tau iHEK. (**D**) Gene Lists of Down-Regulated (Red Boxes) and Up-Regulated (Green Boxes) Genes in WT Tau iHEK. (**E**) Gene Lists of Down-Regulated (Red Boxes) and Up-Regulated (Green Boxes) Genes in P301L Tau iHEK.

Fig S1B displays the scatterplots of WT and P301L tau gene expression, while Fig S1C reports the Principal Component Analysis (PCA). PCA demonstrates significant variation among the study groups. More specifically, the analysis suggests a significant difference in transcriptional activity of WT tau due to the higher number of genes modulated in comparison to the mutant P301L tau form. Using EnrichR^34^, we established Gene Ontology (GO) of the biological processes, molecular functions, and cellular components altered by both the up-regulated and the down-regulated sets of genes. WT tau GO is summarized in Fig 2. WT tau up-regulated genes belonging mainly to classes of cytoskeleton-dependent intracellular transport genes (GO: 0030705, TUBA1A, TUBB2B TUBA1B, TUBB2A and HOOK3) and genes responsible for the regulation of cytoskeleton organization (GO: 0051493). Imbalanced expression of tubulin and tau induces neuronal dysfunction in *C. elegans*,^35^ indicating that tau itself can disturb tubulin gene expression. The reason behind this pronounced involvement of TUBB genes could be due to the fact that *TUBB1B, TUBB2B, TUBA1A* and *TUBB2A* are clustered together within the genome^36^.

**Fig 2.**
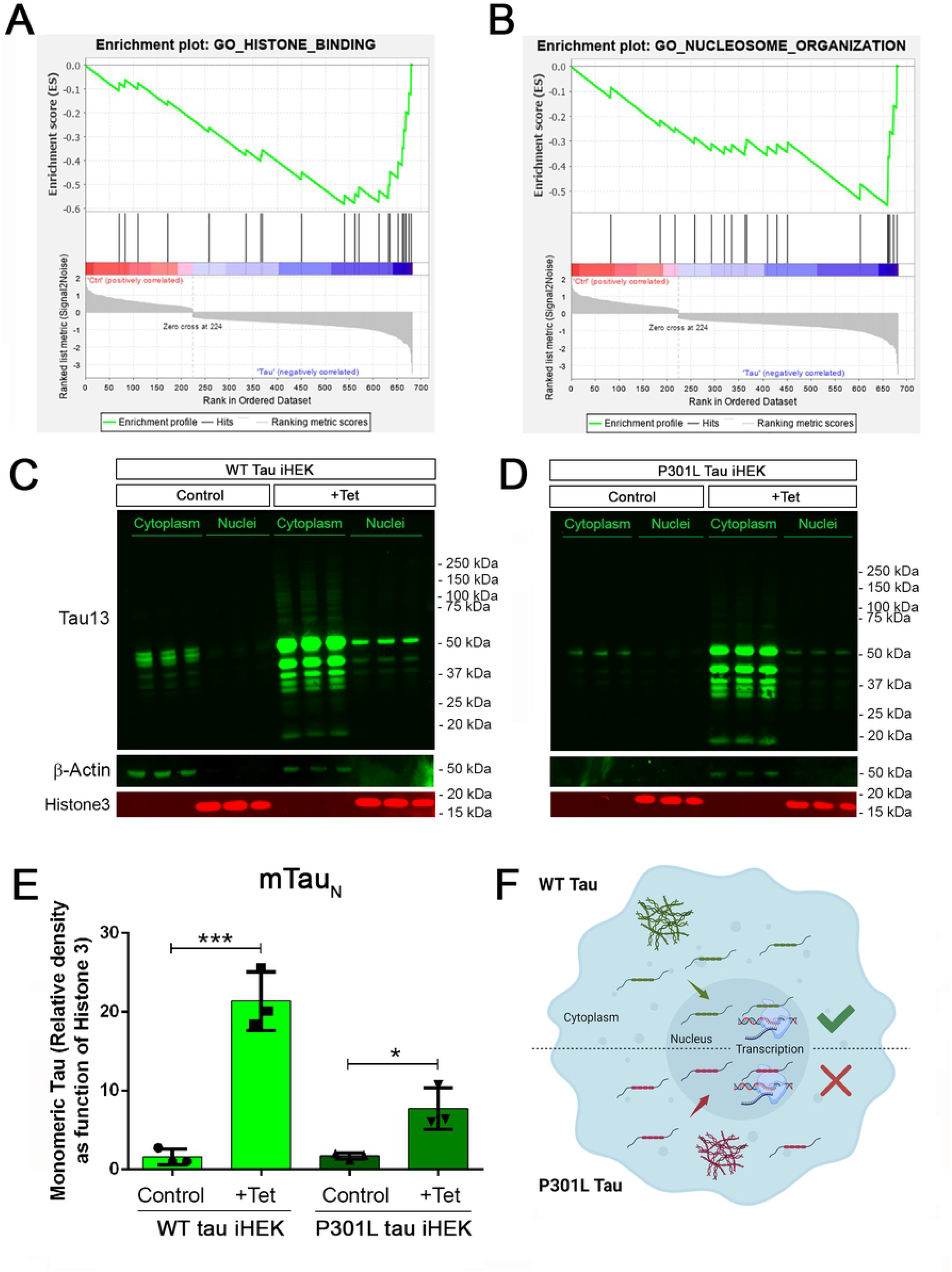
Up- and Down regulated genes in WT Tau iHEK Gene Ontology. Left Column (Green) Up-regulated genes analyzed by Enrich GO and divided by Biological Process, Molecular Function and Cellular Component. Right Column (Blue) Down-regulated genes analyzed by Enrich GO and divided by Biological Process, Molecular Function and Cellular Component. Grey bars represent not significant correlation.

It is important to note that biological process such as microglial cell activation (GO: 0001774) and macrophage activation (GO: 0042116) were also observed as being up-regulated, which confirms known effects of tau on the neuro-inflammatory response commonly observed in neurodegenerative diseases^37^. Neuro-immunomodulation can also effect cytoskeleton reorganization^38^. Our GO analysis revealed up-regulated genes involved in mitochondrion distribution (GO: 0048311, *MAPT* and *MEF2A* genes), morphogenesis (GO: 0070584, *SUPV3L1* p=0.03892), neurogenesis (GO 0022008, *NOM1, MAPT* and *DAGLB* genes), and positive regulation of cell death (GO 0010942, *SAP30BP, MAPT* and *CLU* genes). The increase of *CLU* expression was a particularly interesting observation. Clusterin is a multifunctional, secreted chaperone involved in several basic biological events, including cell death, tumor progression and neurodegeneration. The *CLU* gene is notably associated with an increased AD risk^39^.In terms of molecular functions, the up-regulated genes we observed have several enriched pathways, including RNA binding (GO: 0003723, *USP36, NOM1, TFRC, BAZ2A, SUPTSH, PHF6, FTSJ3, SUPV3L1, TUBA1B, RBM20, MAPT, RBM33, PELP1, HIST1H1C, CPEB4*) and several nuclear functions, such as histone deacetylase binding (GO: 0042826, *MEF2A, SUDS3* and *PHF6*) and sequence-specific double stranded DNA binding (GO: 1990837, *MEF2A, KAT7* and *MAPT*).

Transcriptional products of up-regulated genes are mostly localized in the cytoplasm and nuclear compartments. We detected transcripts associated with nuclear chromatin (GO: 0000790), such as *MEF2A, ZEB2, ANP32E, SUDS3, HIST2H2AC, and HIST1H1C*. We also examined nuclear speck transcripts (GO: 0016607), such as *CARMIL1, USP36, GTF2H2C, BAZ2A, and MAPT* genes, which are also included in nuclear body components (GO: 0016604), along with *SUDS3* and *SENP2*. The other cell compartment well represented in our GO analysis is the cytoplasm. In particular, the microtubule cytoskeleton (GO: 0015630) contained the following up-regulated genes: *TUBB2B, SAP30BP, TUBA1B, TUBA1A, TMOD3, MAP7, TARS, TACC1, MAPT, CLU*, and *RHOQ*. A complete Enrich-GO list of significant up-regulated genes observed in WT tau is presented in Supplemental Table 1.

### WT tau down-regulates genes involved in ubiquitin-related processes as well as genes associated with Golgi and mitochondrial components

Overall, thirty genes were significantly downregulated by WT tau protein. The main biological process affected was the regulation of cellular component organization (GO: 0051128) as it relates to cytoskeleton organization and structure morphogenesis. Molecular functions associated with the aforementioned genes are closely related to ubiquitin protein ligase binding (GO: 0031625) and ubiquitin-like protein ligase binding (GO: 0044389). Genes important to neuronal components included genes essential to the structure of initial axonal segments, nodes of Ranvier, and main axons. These three groups typically involve the gene *KCNQ2*. This gene encodes for Potassium voltage-gated channel subfamily KQT member 2, which plays a critical role in determining the subthreshold electrical excitability of neurons as well as the responsiveness of neurons to synaptic inputs. Therefore, KCNQ2 is important in the regulation of neuronal excitability and the loss-of-function or gain-of function of this gene can lead to various forms of neonatal epilepsy^40^.

Furthermore, Cullin-RING E3 ubiquitin-ligase complex component *KLHL11* is down-regulated, as well as the *STX6* gene. STX6 encodes for Syntaxin-6, which is involved in intracellular vesicle trafficking and is integrally associated with the Golgi apparatus. Another Golgi protein that is down-regulated is Golgin-45 (*BLZF1*). It is required for normal Golgi structure and for protein transport from the Endoplasmic Reticulum (ER) through the Golgi apparatus to the cell surface^41^. Lastly, the ER gene *STC2* is downregulated and encodes for Stanniocalcin-2. This glycoprotein has an anti-hypocalcemic action on calcium and phosphate homeostasis^42^.

We also detected two nucleolus-localized genes among the down-regulated group: UBE2T (ubiquitin-conjugating enzyme with E2 T) and UPF3A, (a regulator of nonsense transcript 3A). The mitochondrial genes that were down-regulated included OXCT1 (Succinyl-CoA: 3-ketoacid coenzyme A transferase 1, mitochondrial enzyme), TRUB1 and PFDN2 (Prefoldin subunit 2). An Enrich-GO list of downregulated genes present in WT tau is depicted in Supplemental Table 2.

Although there are limitations inherent with the model used, these data suggest that WT tau intrinsically and significantly impacts the cell at a transcriptional level. More specifically, a higher number of genes are up-regulated and down-regulated by WT tau when compared to P301L tau. This suggests that the P301L mutation leads to a loss-of-function (LOF) of tau at the transcriptional level. This sort of loss could have detrimental effects on cell structure and organization.

### P301L tau up-regulates gene expression of components related to axonal microtubule skeleton, nuclear speckles, and ribonucleoprotein

The GO pathways and cellular compartments upregulated and downregulated by P301L tau are listed In Supplemental Tables 3 and 4 respectively. As observed in WT tau iHEK cells, the *MAPT* gene is on the upregulated gene list for P301L tau, as expected after Tet induction of the iHEK cells. Within the group of axonal and cytoskeleton genes, we noticed up-regulation of NLGN1, a gene that encodes for Neuroligin-1. Neuroligin is a postsynaptic neuronal surface protein involved in cell-to-cell interactions via its interactions with neurexin family members^43^. It has been established that the NLGN1 gene is associated with amyloid-β oligomers (AβOs) in AD-causing synaptic impairment^44^. In addition, NLGN1 is typically altered in AD hippocampi and also modulates amyloid-beta oligomer toxicity^45^. Neuroligin-1 plays an influential role in synaptic function and synaptic signal transmission, most likely through its ability to recruit and cluster together other synaptic proteins^43^. For instance, neuroligin-1 may promote the initial formation of synapses^46^, but is not essential for the complete formation of synapsyes. *In vitro*, Neuroligin-1 triggers the *de novo* formation of presynaptic structures. NLGN1 may also be involved in specification of excitatory synapses^43^. For example, NLGN1 functions to maintain wakefulness quality and normal synchrony of cerebral cortex activity during wakefulness and sleep^47^. Neuroligin-1 is predominantly located in synaptic cleff of the cell membrane^48^.

When we analyzed upregulated genes, we detected a considerable number of genes related to nuclear body (GO: 0016604) and nuclear speck (GO;0016607) domains including the genes *ITPKC* and *MAPT*. Interestingly, it has been observed that the *FER* gene participates in several different cytoplasmic and nuclear functions. For example, *FER* is associated with nuclear chromatin (GO:0000790) and the microtubule skeleton (GO:0015630). The *FER* gene also encodes for a tyrosine-protein kinase that plays a role in synapse organization, trafficking of synaptic vesicles, the generation of excitatory post-synaptic currents, and neuron-to-neuron synaptic transmission^49^. Lastly, FER plays a role in neuronal cell death after brain damage^49^. The only gene down-regulated by P301L tau is *DCAF12*, which is a component of the Cullin-RING ubiquitin ligase complex^50^. This gene is also down-regulatated by WT tau and belongs to genes associated with ubiquitinization processes. The failure of ubiquitinization pathways is known to have a strong connection to neurodegenrative diseases^51^. Supplemental Figure 2 summerizes upregulated and downregulated genes in P301L tau, subcatagorized by biological process and molecular function.

In summary, the P301L mutation upregulates genes involved in positive regulation of neuronal death and responsivness to reative oxygen species (ROS) production. This is in contrast to the genes altered by WT tau that have a greater affect on cell structural processes. The most important molecular function altered by such genes would be sequence-specific double-stranded DNA binding, transcriptional expression, and chromatin remodelling. Overall, our GO data suggests the precense of both loss-of-function(LOF) and gain-of-function (GOF) events in mutated P301L tau that may relate to pathology. Modulating genes known to be associated with neurodegenerative disease suggests that muatted tau engenders harmful transcription patterns that contribute to the well-established effects of tau proteinaceus-aggregation toxicity.

### WT tau modulates gene expression of chromatin organization and remodeling factors

Gene Set Enrichment Analysis (GSEA) offers an opportunity to evaluate and identify classes of genes or proteins that are over-represented in a large set of genes or proteins and may have an association with disease phenotypes. Due to the differences in gene numbers modulated by WT tau versus P301L tau, we performed GSEA. This analysis compared models with and without WT tau. We observed that WT tau down-regulates the expression of numerous genes linked to chromatin organization (Fig 3A) and chromatin remodeling (Fig 3B) domains. By looking at the chromatin organization and remodeling gene clusters, we identified that several high-mobility group box proteins (HMG) *HMGN5, HMGB2* and *HMGA1* are up-regulated while *HMGB1* and *HMGN1* are down-regulated. It is important to note that HMGB1 is an activator of neuro-inflammatory responses and has been implicated in AD^52^. In addition, several components of the SWI/SFN chromatin remodeling complex are downregulated. The identification of genes *SMARCE1, SMARCA5* and *SMARCC1*, imply that tau has a substantial impact on chromatin remodeling in the cells. The heterogeneous nuclear ribonucleoproteins (hnRNPs), *HNRNPU* and *HNRNPC*, were also found to be downregulated in WT tau. Down-regulation of several factors implicated in DNA replication and repair processes, indicates that WT tau also significantly affects the nuclear compartment of cells in terms of structure and content. Several of these genes are clustered as covalent chromatin modification in GO (Fig 3C).

**Fig 3.**
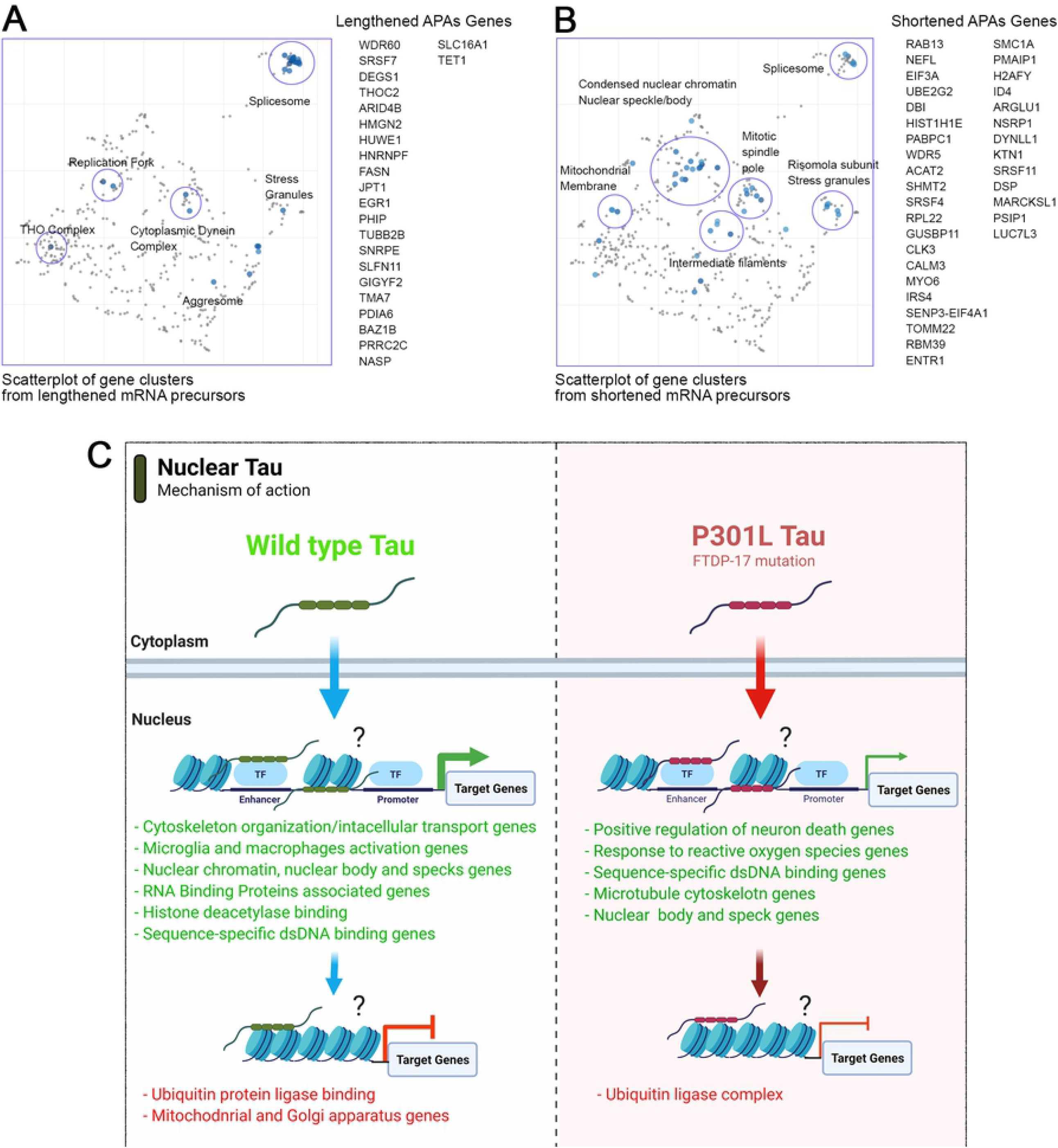
WT tau modulates gene expression of chromatin organization and remodeling factors. (**A**) Enrichment plot for GO Chromatin organization. (**B**) Enrichment plot for GO Chromatin remodeling. (**C**) Enrichment plot for GO-Covalent Chromatin modification. (**D**) IB of Up-regulated genes: RCC1, DNAJC2 and Histone 1.2 (red box) and Down-regulated genes: SMARCC1, SMARCA5 and HMGB1 (blue box) in cytoplasm and nuclear fractions from WT and P301L Tau iHEK. Histone 3 and β-Actin has been used as loading control for nuclear and cytoplasmatic fractions, respectively. (**E**) representative Tau 13 (magenta) and Histone 1.2 (green) Co-IF of control (-Tet) and treated WT Tau iHEK. (**F**) Histone 1.2 integrated density quantification in control and +Tet cells (Unpaired t-test, *p<0*.*0001*, ****). (**G**) representative Tau 13 (magenta) and Ki67 (green) Co-IF of control (-Tet) and treated WT Tau iHEK. (**H**) Ki67 integrated density quantification in control and +Tet cells (Unpaired t-test, *p<0*.*0001*,****). (**I**) representative Tau 13 (magenta) and SMARCC1 (green) Co-IF of control (-Tet) and treated WT Tau iHEK. (**J**) SMARCC1 integrated density quantification in control and +Tet cells (Unpaired t-test, *p<0*.*0001*,****). (**K**) representative Tau 13 (magenta) and SMARCA5 (green) Co-IF of control (-Tet) and treated WT Tau iHEK. (**L**) SMARCA5 integrated density quantification in control and +Tet cells (Unpaired t-test, *p<0*.*0001*,****).

To validate gene expression changes observed in GSEA analysis, we verified multiple proteins via western blot by using the up-regulated and down-regulated lists generated from Histone Binding GO. We verified up-regulation of RCC1, DnaJC2 and Histone1.2 proteins in the cytoplasm and in nuclear fractions of WT and P301L tau iHEK cells (Fig 3D). We also confirmed RCC1 expression and noticed its accumulation in the cytoplasm for both cell lines. Interestingly, we discerned that RCC1 is not imported into the nuclei where it should function as a regulator of chromatin condensation. Instead, DnaJC2 in P301L tau iHEK cells appear to be downregulated. However, Histone 1.2 is upregulated in both cell lines. We did not observe down-regulation of the chromatin remodeling complex factors SMARCC1 and SMARCA5. Instead, we detected their accumulation in the cytoplasmic fractions while in the presence of tau, which suggests a deficit in these factors in the nuclei, as observed in our western blots. Lastly, HMGB1 and β-Actin are down-regulated, but HMGB1 is not detected in the nuclei when in the presence of tau. Histone 3 was used as a nuclear loading control.

To verify gene expression results, alongside western blots, we performed co-immunofluorescence in WT tau iHEK cells. We evaluated integrated density of Histone 1.2 (Fig 3E and 3F), Ki67 (Fig 3G and 3H), SMARCC1 (Fig 3I and 3J), and SMARCA5 (Fig 3K and 3L). Analysis was performed by considering nuclear integrated density of “–” and “+” tau WT iHEK proteins. To detect and confirm tau expression, we used the Tau13 antibody. MCM2 and RCC1 images and their relative integrated density quantifications are presented in Supplemental Fig. 4. GSEA analysis for WT tau revealed significant down-regulation in the pathways for histone-binding (Fig 4A) and nucleosome organization clusters (Fig 4B). Several genes were detected in the histone and nucleosome domains, which were recurring and can be viewed in the chromatin gene list showed in Fig 3. In addition, we observed an up-regulation of *RCC1* (a regulator of chromosome condensation), *CTSL* (Cathepsin L), *MCM2* (Minichromosome maintenance complex component 2), and *DNAJC2* (DnaJ heat shock protein member C2). In Nucleosome GO, we observed up-regulated *HMGB2* and *HMGA1* (high mobility group box B2 and A1). On the contrary, several Lysine acetylation regulators were downregulated: *BRD3* and *BRD9* (from BRD family), *HDAC2, KDM5B, KAT7, and SFTD2*.

**Fig 4.**
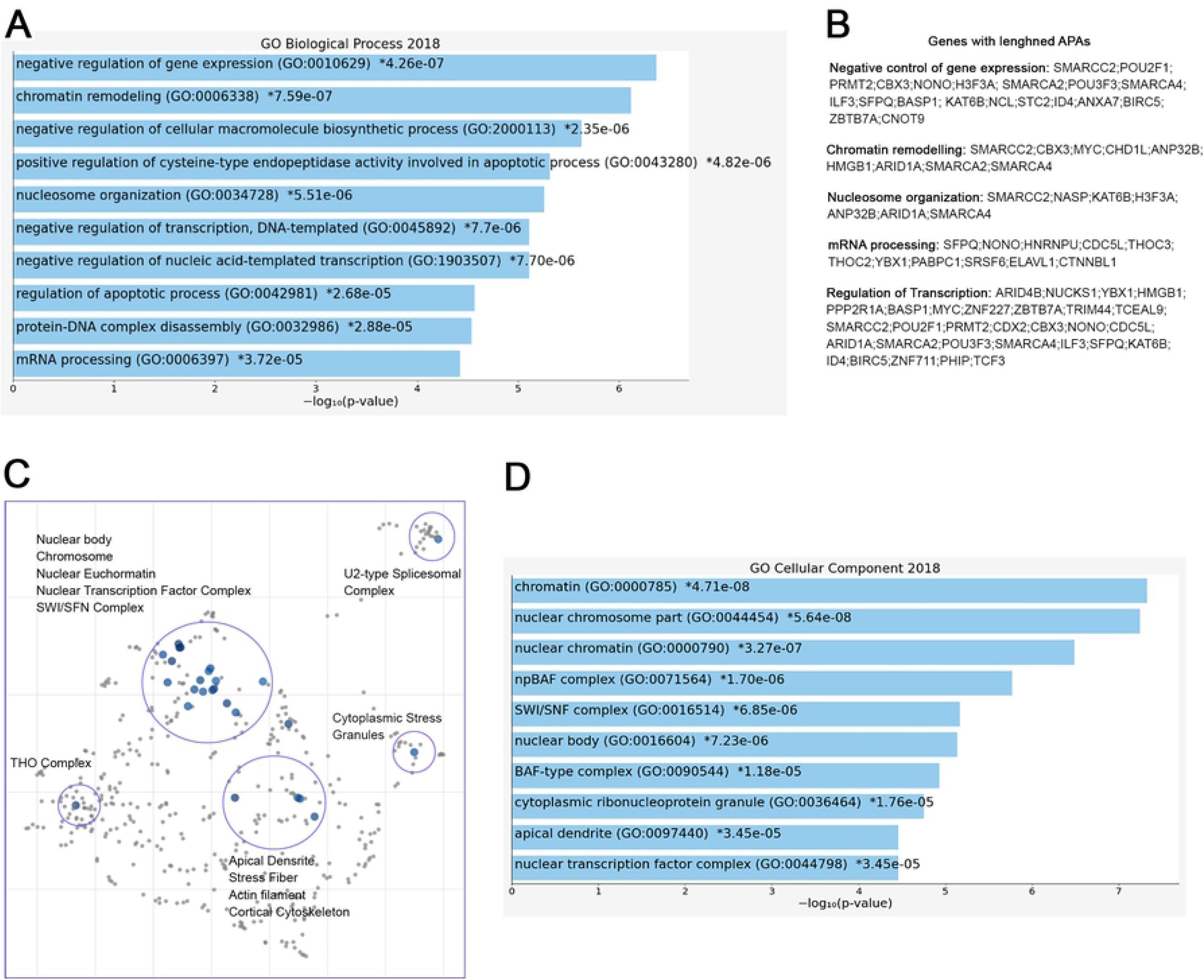
Tau nuclear shuttling. (**A**) GWAS GO-Histone Binding heat map in WT Tau. (**B**) GWAS GO-Nucleosome organization heat map in WT Tau. (**C**) Enrichment plot for GO Histone Binding. (**D**) Enrichment plot for GO Nucleosome Organization. (**E**) Immunoblot (IB) with Tau13 (1:1000) and β-Actin () of cytoplasm and nuclear fraction from WT (left panel) and P301L (right panel) Tau induced with Tet. (**F**) Relative density of nuclear monomeric Tau (mTau_N_, normalized with Histone3). Unpaired *t-test* has been performed to compare column means ((-) WT Tau vs WT Tau ***, *p=0*.*0009*, (-) p301l Tau vs P301L Tau *, *p=0*.*0169*, WT Tau vs P301L Tau **, *p=0*.*0065*). (**G**) Schematic model on Tau nuclear import in the two iHEK cell lines.

We also used western blotting to verify tau levels in cytoplasm and nuclear fractions of WT and P301L tau iHEK cells (Fig 4C and 4D, respectively). We found that upon Tet induction in both compartments, tau was detected, which was previously observed^7^ and expected. Western blot analysis demonstrated that tau is represented mainly in its monomeric form (mTau_N_) when probing the nucleus. We compared the level of mTau_N_ in both cell lines and we determined that mTau_N_ increased in both cell lines after Tet induction. However, the WT mTau_N_ was present in a significantly higher level when compared to the P301L mTau_N_ (Fig 4E). This difference is due to the higher *MAPT* transgene expression efficiency in WT tau iHEK cell lines as was confirmed by RT-qPCR in a previous study^7^. These observations suggest that the monomeric form of tau protein predominantly carries out transcriptional activity and that the P301L mutation did not affect the nuclear import of tau, but instead modulated transcriptional activity. Cytoplasmic mTau was quantified as well (Supplemental Figure 4). In general, we propose that WT and P301L tau both shuttle into the nuclei but then modulate transcription differently. The schematic model for this idea is represented in Fig 4F. In summary, many nuclear factor genes involved in several nuclear activities, including chromatin condensation, are downregulated in WT tau, which indicates a potential role of WT and P301L tau in the control of chromatin factors, expression and subsequent cellular localization.

### RNA metabolism, chromatin organization and *HNRNP*s precursor’s display shortened APAs in the presence of WT tau

From PAC-seq analysis, we identified 110 genes with shortened 3’UTRs. The majority of these shortened genes belong to significant pathways associated with mRNA processing (GO: 0006397), RNA Splicing (GO: 0000377, GO: 0000398), and RNA metabolic processes (GO: 0016070) (Fig 5A). These domains share several genes: *HNRNPA3, SRRT, PRPF4B, CCAR1, LSM8; SNRNP40, HNRNPK, ZMAT2, ZC3H11A, HNRNPF, PCBP2, SNRPE*, and *HNRNPC*. The regulation of responses to DNA damage (GO: 2001020) comprise the following genes: *BCLAF1, FMR1, USP1* and *HMGA2* (others listed in Fig 5B).

**Fig 5.**
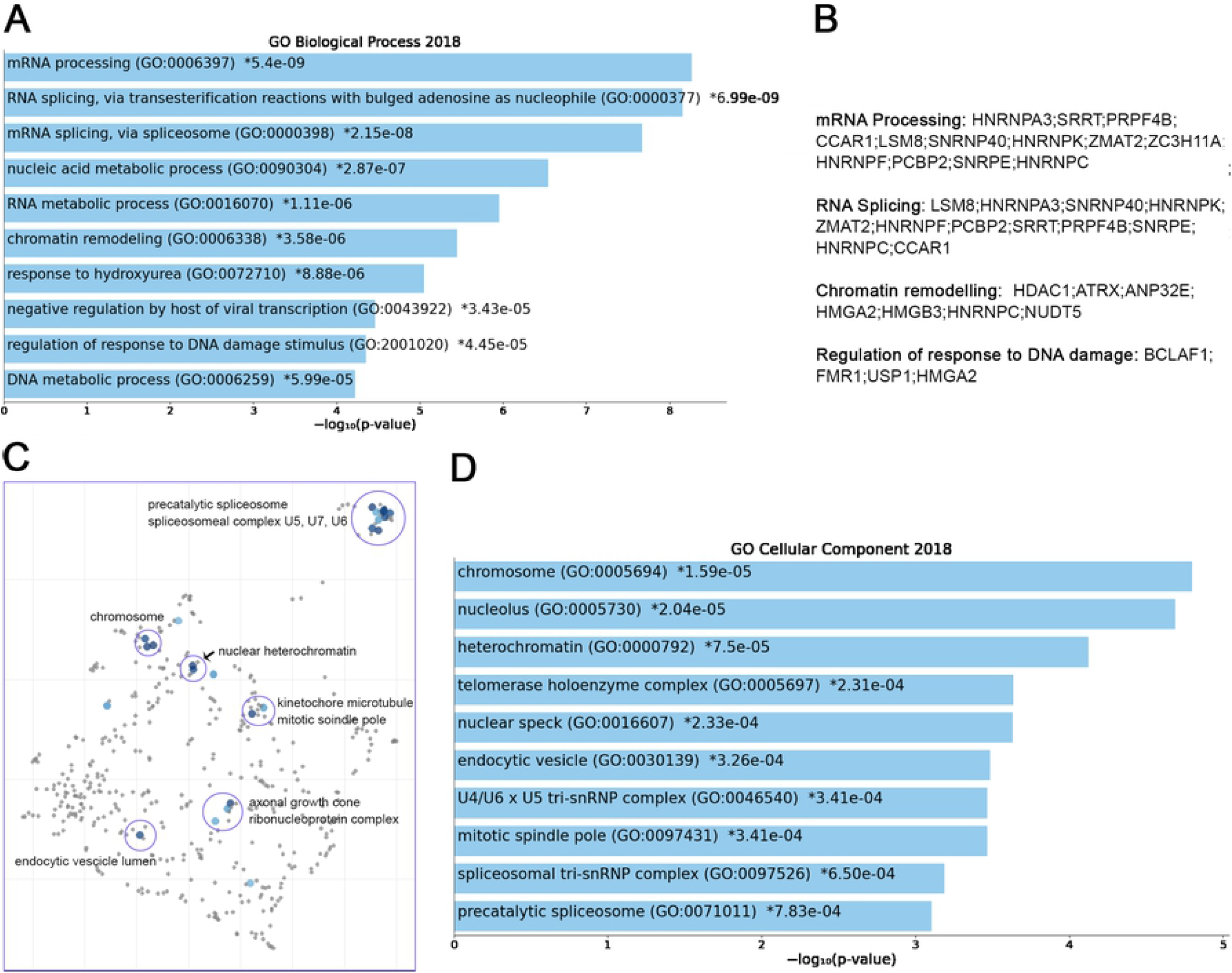
RNA metabolism, chromatin organization and *HNRNP*s precursor’s display shortened APAs in the presence of WT tau. (**A**) Enrich-GO Biological Process of WT Tau shortened APAs (p-value reported). (**B**) Partial list of biological process genes with shortened APAs upon presence of WT Tau (mRNA processing, RNA splicing, chromatin remodeling and regulation of response to DNA damage). (**C**) The Cellular Components scatterplot is organized so that similar gene sets are clustered together. The larger blue points represent significantly enriched terms - the darker the blue, the more significant the term and the smaller the p-value. The gray points are not significant. Plots has been generated and downloaded using scatter plot visualization Appyter. (**D**) Enrich-GO Cellular Component of WT Tau shortened APAs (p-value reported).

Within the shortened APA precursors, various genes are related to nuclear function, such as the chromosome related genes (GO: 0005694) *IK, FMR1, HMGA2, SMC4, SMC3, SMC2* and *SMC6*. Structural maintenance of chromosome (SMC) proteins are ATPases that are essential to chromosomal condensation, sister-chromatid cohesion, recombination, DNA repair, and epigenetic silencing of gene expression^53^. Eukaryotes have at least six genes encoding SMCs (SMC1-SMC6)^54^. They inherently work as heterodimers: SMC1/SMC3 (Cohesin Complex), SMC2/SMC4 (Condensin Complex) and SMC5/SMC6^54^.

Several nucleolar (GO: 0005730) genes have altered poly(A) site usage by WT tau including: *PARP1, FMR1, CHD7, DDX21, PWP1, PPM1E, SMC2, RSL1D1, ILF3, NCL, S100A13, KIF20B, RAN*, and *GET4*. As we saw in the shortened 3’UTRs, the most affected genes for lengthened 3’ UTRs lie within the RNA binding function domain (GO: 0003723). MRNA processing, RNA splicing, and nucleic acid metabolic processes received the top scores, indicating a strong impact of WT tau in the regulation of mRNA isoforms at different levels. All significant enrichment terms are clustered and represented in a scatterplot in Fig 5C. In the mRNA processing domain (GO: 0006397) we identified several heterogeneous nuclear ribonucleoproteins (hnRNPs) genes (*HNRNPA3, HNRNPK, HNRNPF, HNRNPC, HNRNPDL*). HnRNPs are involved in alternative splicing, transcriptional and translational regulation, stress granules formation, cell cycle regulation, and axonal transport^55^. Their dysfunction has been shown have neurological implications, but their roles have not been comprehensively investigated. Several neurodegenerative diseases, including AD, FTD, and amyotrophic lateral sclerosis (ALS) have been associated with hnRNPs when it comes to the progression of these pathologies^56^. More specifically, hnRNPK has been linked to the transcripts of several cytoskeletal genes, including *MAPT*, which is needed for axonogenesis^57^.

In Alzheimer’s disease, hnRNPC promotes APP translation^58^ and stabilizes the APP precursors mRNA, which could suggest that increasing hnRNPC levels may promote Aβ secretion^59^. Within the hnRNPs group, hnRNPA3, hnRNPF and hnRNPDL are all detected in pathological inclusions of ALS and FTD brains^56,60,61^. Moreover, hnRNPK is a regulator of p53^62^,which we and others recently discovered was present in elevated amounts in AD cortices^11,12^. It has been also determined that hnRNPK sumoylation mediates p53 activity^63^. All this evidence places hnRNPs in a central position for further experimental analysis in human brain tissues to elucidate more valuable information about the localization and function of this large family of ribonucleoproteins.

HnRNPA3 has been identified in neuronal cytoplasmic and intranuclear inclusions in patients with GGGGCC expansion repeats^61^ and hnRNP F were also found to co-localize with GGGGCC expansion foci in immunoprecipitation studies^64^. In addition, western blot analyses imply that hnRNP may be in part responsible for the toxicity incurring by C9orf72 mutations, considering important RNA processes such as splicing are compromised. hnRNP A3 and K have been found associated with TDP-43^65^. Implications of tau-mediated APAs in hnRNPs open new venues for investigators to study new mechanistic insights of these proteins in several proteinophaties. Within RBPs group, we also observed the *MATR3* gene. This gene encodes for Matrin3, a DNA/RNA-binding protein. Mutations in this gene cause familial ALS/FTD, and MATR3 pathology is a feature of sporadic disease, suggesting that its dysfunction is inherently linked to ALS pathogenesis^66^.

Shorter 3’UTR are generally associated with enhanced translation of the mRNA APA in the presence of WT tau, which supports the finding that high-levels of hnRNPs sustain dysfunction of stress granules in ALS and FTD. Recent proteomic analysis in AD human Neurofibrillary Tangles (NFTs) showed that phospho-tau in NFTs is associated with more than 500 proteins^67^. We observed several of these proteins in the APAs shortened WT tau, such as HNRNPK, ILF3, AP2B1, RAN, RAB11A, HSP90B1, PARP1, MATR3, PPIA, NCL, HNRNPA3, HSP90AA1, and HNRNPC. It is intriguing that the presence of chaperone Hsp90, a tau-regulated gene, plays a crucial role in neurodegenerative pathologies and has been studied in AD or a long time^68^.

These observations suggest that tau has early effects on gene expression that results in later stages of toxic associations commonly found in neurodegeneration. Enrich-GO (Cellular Function) of shortened-APAs genes by WT tau is provided in the supplemental information section. GO-Cellular Process, Molecular Process and Cellular Components bar charts of shortened APAs are shown in Fig S4.

### SWI/SFN, THO complexes, and several RNA-Binding protein precursors display lengthened 3’UTRs in presence of WT tau

Further analysis revealed 173 genes with lengthened APAs. The complete list of the 173 genes with lengthened APAs is reported in the supplemental information section. Among these genes, we found that many of them are related to three major biological process: chromatin remodeling (GO:0006338), negative regulation of gene expression (GO: 0010629), and mRNA processing (GO:0006397) (Fig 6A). To be more specific, we noticed several genes belonging to the ATP-dependent chromatin remodeling complex npBAF (mammalian SWI/SFN, GO: 0071564): *SMARCC2, ARID1A, SMARCA2* and *SMARCA4*. This complex is found in neuronal progenitor’s cells and post-mitotic neurons, and it is essential for the maturation of the post-mitotic neuronal phenotype as well as long-term memory formation^69^. Along with the chromatin remodeling complex, other genes contained altered APAs, including pericentric chromatin components (GO: 0005721, *HELLS* and *CBX3*), and nuclear chromatin factors (GO:0000790, *SMARCC2, CBX3, H3F3A, NUCKS1, ARID1A, SMARCA2, SMARCA4, HIST2H2AC, RAD50, NASP, MYC, NSMF, TCF3*) (Fig 6B).

**Fig 6.**
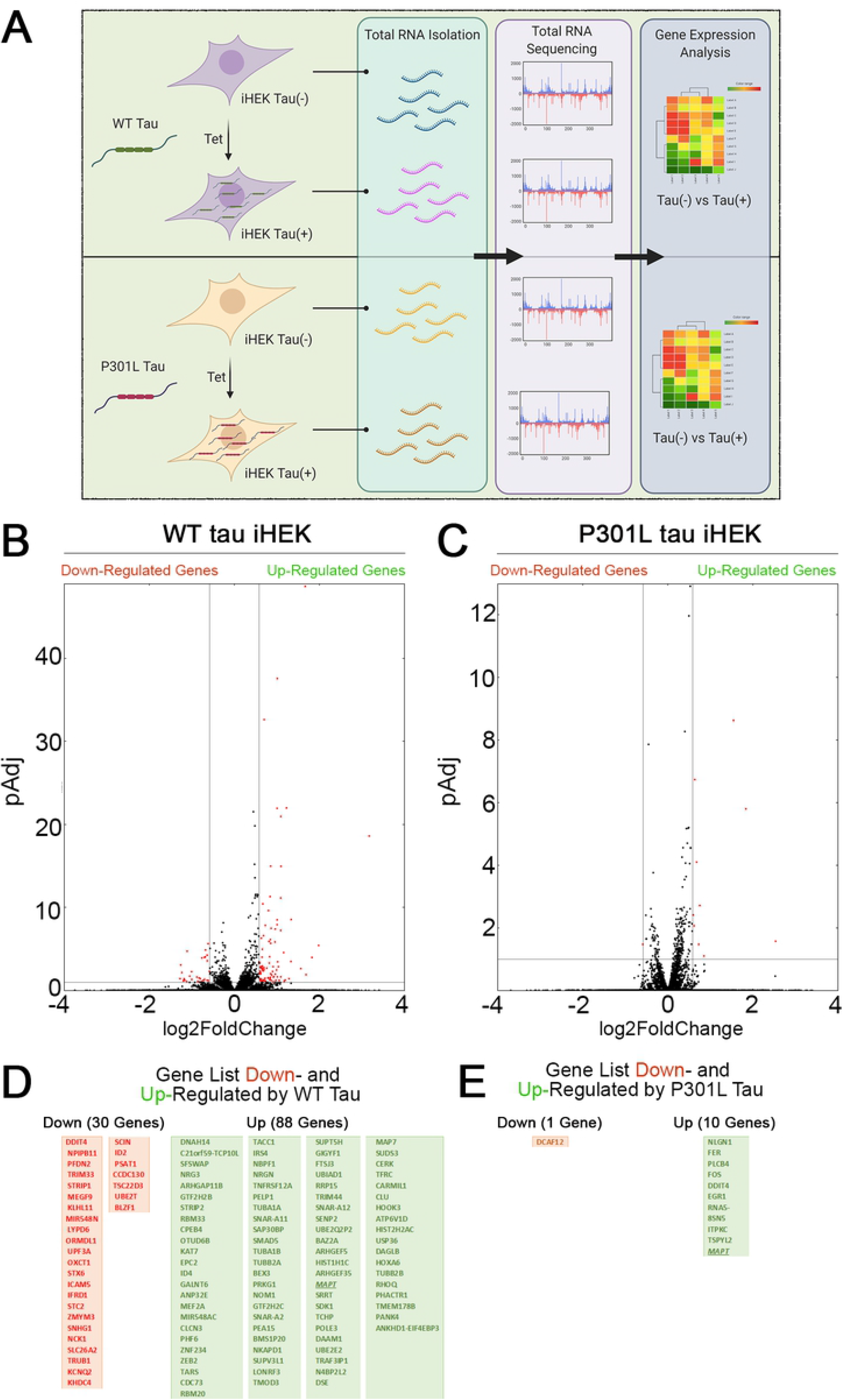
RNA processing and splicing precursor’s display lengthened APAs in presence of WT tau. (**A**) Enrich-GO Biological Process of WT Tau lengthened APAs (p-value reported). (**B**) Partial list of biological process genes with lengthened APAs upon presence of WT Tau negative control of gene expression, chromatin remodeling, nucleosome organization, mRNA processing and regulation of transcription). (**C**) The Cellular Components scatterplot of lengthened APAs in WT Tau is organized so that similar gene sets are clustered together. The larger blue points represent significantly enriched terms - the darker the blue, the more significant the term and the smaller the p-value. The gray points are not significant. Plots has been generated and downloaded using scatter plot visualization Appyter. (**d**) Enrich-GO Cellular Component of WT Tau lengthened APAs (p-value reported).

Several nuclear speck (GO: 0016607) genes were also identified: *BASP1, POM*,; *ERBI, YLPM1, HNRNPU, LUC7L3, CDC5L, TCF3, SRSF6*, and *KIF20B*. Cytoplasmic ribonucleoprotein granule (GO: 0036464) and cytoplasmic stress granules (GO: 0010494) genes were delineated as *MBNL1, CARHSP1, NCL, HNRNPU, IQGAP1, YBX1, RAC1, PABPC1, CNOT9*. Within the domain of RNA processing, two genes *THOC2* and *THOC3* were also identified. They are components of the THO complex (GO: 0000445) involved in efficient export of poly-adenylated RNA and spliced RNAs^25^.

The THO complex appears to coordinate transcripts for synapses development and dopamine neuron survival^70^. Recently, it has been found to interact with ZC3H14, which regulates the processing of neuronal transcripts^71^, so it is not surprising to find in our dataset another polyadenosine RNA-binding protein *ZC3H15* on the list of lengthened APAs. These observations indicate that export complex RNA precursors are meaningfully affected by WT tau.

Not surprisingly, many translation initiation factors (GO: 0003743) were also discovered in our analysis including *EIF2S3, EIF3E, EIF3A, EIF1*, and *EIF4G1*. It is important to note that many APA-lengthened proteins in our study are RNA-Binding Proteins (RBPs). In fact, 46/173, or ∼27% of the total were. RBPs are implicated in the pathogenesis and progression of numerous neurodegenerative diseases, and they are linked to toxic interactions and aggregations in amyloidogenic proteins such Amyloid-beta and tau. The subsequent dysfunction of RBPs is closely related to distinct pathways that are altered in proteinophaties^72^.

Considering the above, we also studied the presence of lengthened APAs of *ELAVL1*. This gene encodes for HuR (RBPs), which is a neuroprotective protein. This protein has been demonstrated in the regulation of oxidative metabolism in neurons as a way to protect from neurodegeneration^73^.

Apical dendrites (GO: 0097440) (*MAP1B, NSMF and CLU*) and other cytoskeletal genes (*ACTR2, LIMA1, TPM4, PPP2R1A, BASP1, TARS, PHIP, NSMF, IQGAP1, RAC1, CLU*, and *SMARCA2*) display lengthened poly-A tails as well. Enrich-GO (Cellular Function) of lengthened-APAs genes in WT tau is provided in the supplemental information section. GO-Cellular Process, Molecular Process and Cellular Components bar charts of lengthened-APAs are shown in Fig S4.

### P301L tau modulates 3’UTRs of RNA export complex THOC and splicing precursors SNRPE

In P301L tau precursor APAs, we detected 23 lengthened genes in total. More specifically, the THOC2 gene, which is a component of the THO complex (GO: 0000445) was lengthened in WT tau. Another gene of the small nuclear ribonucleoprotein complex (SNRPE) was detected. *SNRPE* is also a gene for the spliceosome complex (GO: 0005681) (Fig 7A). Lastly, the nuclear replication fork (GO: 0043596) gene *BAZ1B* was also observed.

**Fig 7.**
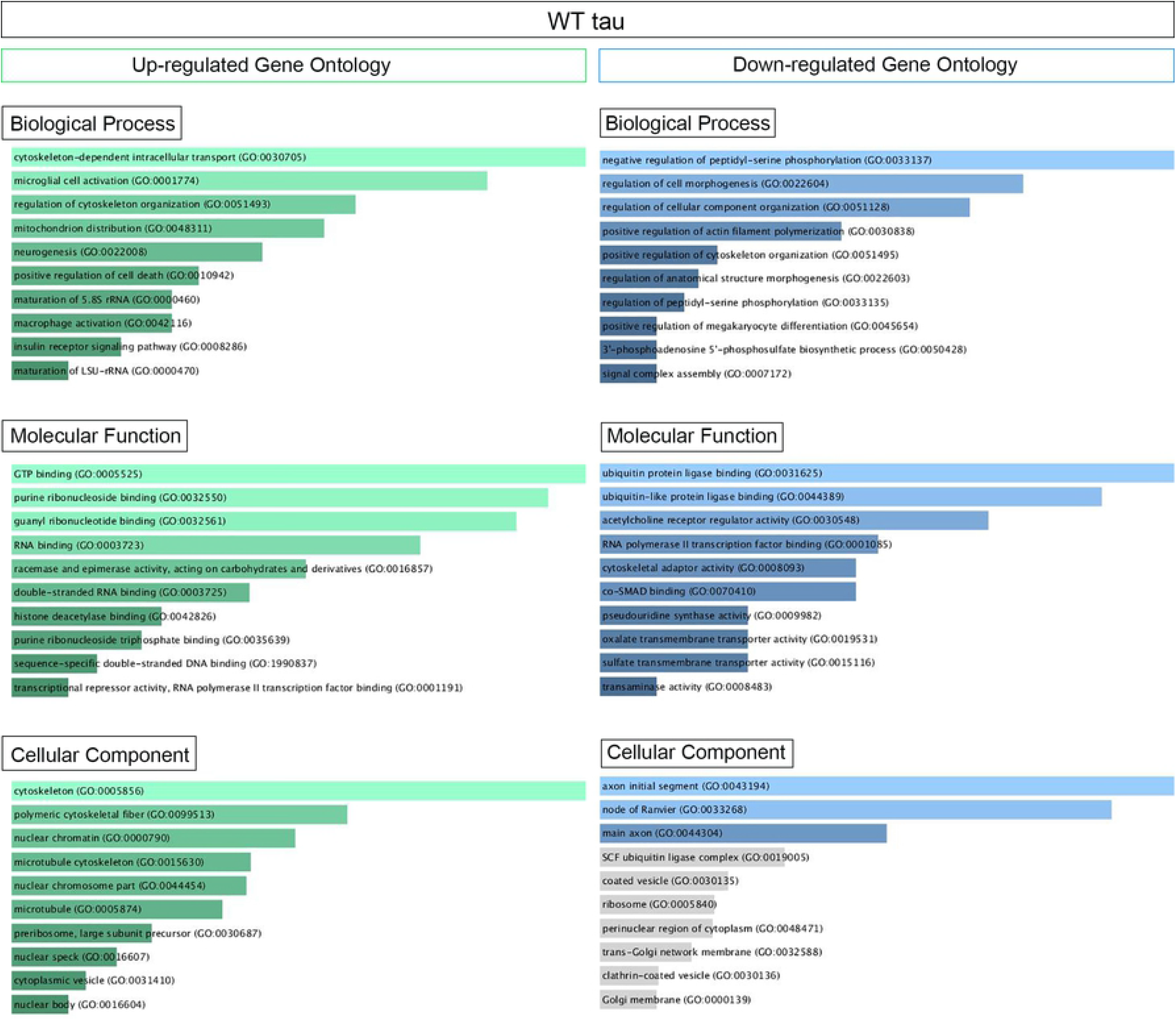
Mutant P301L Tau modulates APAs associated with spliceosome and nuclear chromatin. (**A**) Scatterplot of gene clusters from lengthened mRNA precursors upon P301L Tau expression. (**B**) Scatterplot of gene clusters from shortened mRNA precursors upon P301L Tau expression. (**C**) Model for nuclear Tau activity to transcriptional and post-transcriptional levels.

In contrast to WT tau, P301L tau induces lengthening of the *HNRNPF* gene. HnRNPs represent a large RNA-Binding protein family that contributes to many aspects of nucleic acid metabolism, including alternative splicing, mRNA stabilization, transcriptional, and translational regulation^55^. Dysregulation of RNA metabolism is crucial in the pathogenesis of several neurodegenerative diseases as Parkinson’s^74^, FTD and overlaps with aspects of ALS. Some studies revealed possible involvement of hnRNPs in the pathogenesis and progression of these diseases^75^. Furthermore, hnRNP F has been uncovered in RNA foci in human brain tissue of FTD-ALS patients^56^. Affinity pull-down assays and genome-wide analysis also revealed a hnRNP F-bound splicing complex that regulates neuronal and oligodendroglial differentiation pathways in the developing brain^64^. As observed for WT tau, the mutant P301L form also modulates several RNA-Binding Proteins (GO: 0003723): *SLFN11, HNRNPF, FASN, HUWE1, PRRC2C, THOC2, HMGN2, SRSF7*, and *GIGYF7*. We found 34 genes in total with evidence of APA and shortened 3’UTRs (Fig 7B). The three top-scored cellular components were nuclear speck (GO: 0016607), nuclear body (GO: 0016604) with *RBM39* (ALS associated gene^76^) and Nuclear heterochromatin genes (GO: 0005720). Nuclear speak and body genes consisted of *LUC7L3, SRSF4, NSRP1 and SRSF11*. Nuclear heterochromatin genes detected were *H2AFY* and *HIST1H1E. H2AFY* encodes for a variant of the H2A histone that is present in a subset of nucleosomes where its role is to represses transcription^77^.

The Cellular Components scatterplot of lengthened APAs in WT Tau is presented in Fig 6C and GO Cellular component bar charts in Fig 6D.

These data suggest that the mutant P301L form of tau reduces activity in transcription and alternative poly(A) tails processes due to loss-of-function. However, P301L tau does generate different mRNA isoforms of transcripts mainly translated in splicing factors, nuclear speckle/body structures and chromatin remodeling proteins. Enrich-GO (Cellular Function) of shortened and lengthened-APAs by P301L tau is provided in the supplemental information section. GO-Cellular Process, Molecular Process and Cellular Components bar charts of shortened and lengthened-APAs are shown in Fig S4.

## Discussion

In this study, we revealed new mechanistic insights into non-canonical tau functions. In particular, we showed novel tau activities in transcription and alternative poly-adenylation (APA) pathways. APA is a widespread mechanism of gene regulation that generates 3’ ends in transcripts made by RNA polymerase II^78^. APA is regulated in cell proliferation, differentiation and extracellular cues. It occurs in the 3’UTR and leads to the production of mRNA isoforms, followed by splicing which leads to the production of distinct protein isoforms^78^. Tau is typically described as an abundant neuronal microtubule-binding protein. Recently, we observed its presence within non-neuronal human cell lines and neuronal nuclei in AD brains ^7,13^ alongside other study^2^. We were particularly interested in the possibility of non-canonical tau functions. We hypothesized that nuclear tau acts as a transcriptional regulator. To test our hypothesis, we used the tau inducible HEK system, which is a well-established cell line capable of studying mechanisms related to the tau aggregation process within a controlled system of MAPT gene expression^79^. Our study employed new technologies such as Poly(A)-ClickSeq to resolve whether genes were upregulated or downregulated by WT and P301L tau in an *in-vitro* model. Furthermore, we analyzed alternative polyadenylation (APA) profiles under the presence of WT and P301L tau^76^.

Our results suggest that both WT and P301L tau are able to shuttle into the nuclei (Fig 4). This observation confirmed our previous observations ^7^. We did not investigate the effect of the P301L mutation on nuclei-cytoplasm shuttling in this report. The decreased number of genes expressed in P301L cells suggests that this particular mutation of tau impairs transcriptional activity. We did not investigate the LOF consequences of P301L tau in great detail, but our observations suggest new mechanistic insights linked to alternative nuclear tau function.

One APA transcript of significance is the *SFPQ* gene, which we identified in WT tau expression as having a lengthened 3’UTR. *SFPQ* has been associated with tau as a critical factor for rapid progression of AD, and it has been observed as downregulated in post-mortem brain tissue of rapidly progressive AD patients^80^. Therefore, the lengthened APAs in this gene could explain the down-regulation in the presence of a high level of tau, which mimics late-stage AD. In-vitro data of SFPQ down-regulation due to human tau suggest a causal role of tau, possibly through the alternative poly-adenylation of *SFPQ* transcripts.

Further analysis comparing 3’UTRs lengthened between WT and P301L tau revealed that a significant number of RBPs showed lengthened 3’UTRs in P301L compared to WT tau. For example, we detected 72 RBPs including *FUS* (found in the supplemental information section). These data suggest a significant difference in RNA isoforms based on genetic tau background, which then subsequently modulates different aspects of RNA metabolism in neurons.

Using the same cellular models, we determined that the prominent form of nuclear tau is monomeric, but Tet induction causes tau oligomerization within the nuclei^7^. The formation of large and nuclear oligomeric forms is another possible explanation for LOF observed as a consequence of mutated tau. Mutant P301L tau shows a distinct aggregation mechanism compared to WT^81^ and aggregates faster than WT^82,83^. For example, monomeric tau in the cytoplasm of cells producing (WT or P301L) tau aggregate and subsequently avoid nuclear translocation. In addition, aggregation in the cytoplasm and within the nuclei of tau reduces the pool of monomeric nuclear tau. This pathological mechanism can compete with functional monomeric and oligomeric tau, which then alters tau transcriptional activity. This phenomenon should be investigated in the near future using neuronal models. Another function of tau is binding DNA in-vitro. Overall, the multifunctional nature of nuclear tau should be thoroughly scrutinized in order to identify unrevealed functions connected to DNA expression and RNA processing. We suggest that the nature of nuclear tau as a transcriptional factor, chromatin remodeler and/or transcriptional co-factor must be elucidated using proper models such as induced pluripotent stem cells or mouse primary neurons carrying mutation on P301 site. At this stage, we can only hypothesize the direct and indirect effects of tau during transcription.

This study utilized PAC-ClickSeq technology to identify the APA modulated by P301L and WT tau. Alternative Poly-A (APA) sites in human genome have been identify mainly in 3’UTRs (UTR-APA) sites, which harbor diverse regulatory sequences. This type of APA can change the length and composition of 3’UTR, which subsequently affects the binding of miRNAs and/or RBPs. This post-transcriptional modification leads to differences in mRNA stability, export, localization, translational efficiency^26^. Although the currently accepted theory is that genes with longer 3’UTR tend to show decreased expression levels, this does not necessarily mean that every single gene with a longer 3’UTR is less stable those with a shorter one.

We plan to investigate these findings using primary neurons and in-vivo models in the near future. We are choosing these alternative models because the iHEK cell model have inherent limitations in terms of reliability as a neuronal system. However, the iHEK cells used in this study are an established model used by many researchers to study the mechanistic insights of tau aggregation and toxicity. The results presented in this study support non-canonical functions of tau. Therefore, we report broad tau-driven, post-transcriptional regulation in APAs by both WT and P301L tau considering both cell lines produced high levels of monomeric and aggregated tau. In this study, we did not investigate which tau isoform regulates APA in cells and by what method tau regulates APAs, but we established a new category of interest in post-translational modification. We hope further studies of nuclear tau and its relation to DNA and RNA processing will identify new targets in tauopathies and eventually find new therapeutic targets.

## Limitations of the study

As mentioned in the discussion, the main limitation of this study is the nature of tau inducible HEK cells. We are aware that further study on neuronal cells is necessary. However, iHEK models are commonly used to study mechanisms that are tau-dependent and several of them have been translated into neurons models. All relevant datasets used and/or analyzed in this current study are available upon request from the corresponding author.

## Supplemental Information

The source data underlying all main and supplementary figures are provided as a Source Data file. RNAseq datasets is uploaded to NCBI SRA, reference number: <u>PRJNA744518</u>. Figure 1A, 4F and 7C were generated using BioRender Software (https://biorender.com).

## Acknowledgments and Funding

We thank the members of the Kayed and Routh labs for their support and help. We thank Bergman Isabelle B. and Leiana Fung for editing and proofreading of the manuscript. This work was supported by Mitchell Center for Neurodegenerative Diseases, the Gillson Longenbaugh Foundation and National Institute of Health grants: R01AG054025, R01NS094557, R01AG055771, R01AG060718 and the American Heart Association collaborative grant 17CSA33620007 (R.K.).

## Author contribution

Conceptualization, M.M., A.R. and R.K.; Methodology, M.M., A.R. and R.K.; Investigation, M.M., E.J., S.M., A.E. and S.G.; Transcriptomic analysis, A.R. and E.J.; Writing – Original Draft, M.M.; Writing – Review & Editing, all authors; Funding Acquisition, R.K.; Resources, R.K.; Supervision, M.M. and R.K.

## Abbreviations

AD: Alzheimer’s disease
APA: alternative polyadenylation
DDR: DNA damage response
FTD: frontal temporal dementia
GO: Gene Ontology
GSEA: gene set enrichment analysis
GWAS: genome wide association study
MTs: microtubules
PAC: poly(A) cluster
PAC-Seq: Poly(A)-ClickSeq
PAP: poly-A polymerase
PAS: poly(A) site
RNA: ribonucleotide acid
Tet: Tetracycline
3’UTR: 3’ untranslated region
iHEK: inducible human embryonic kidney cells
ER: endoplasmic reticulum

## Figure Legends

**Supplemental Table 1.**
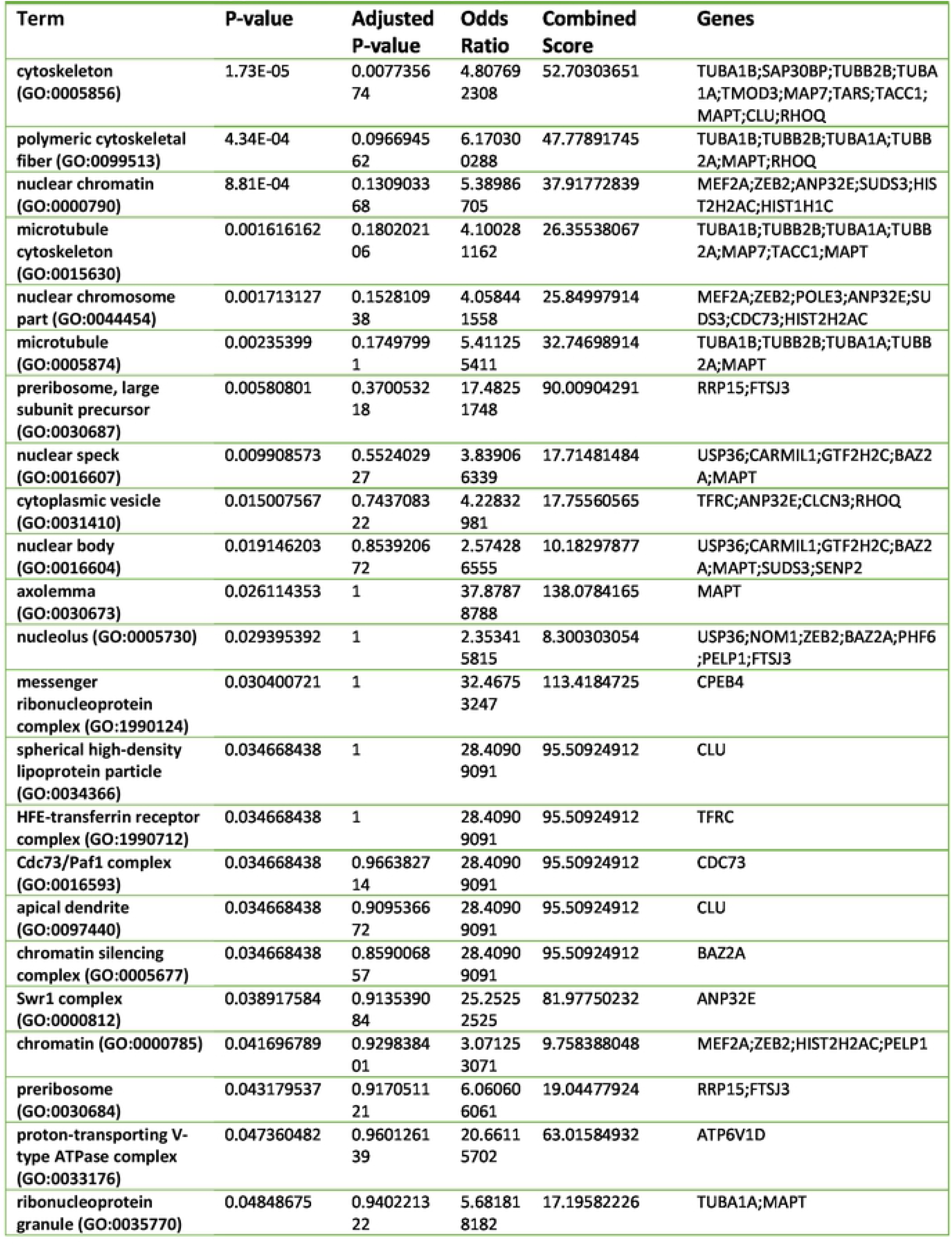
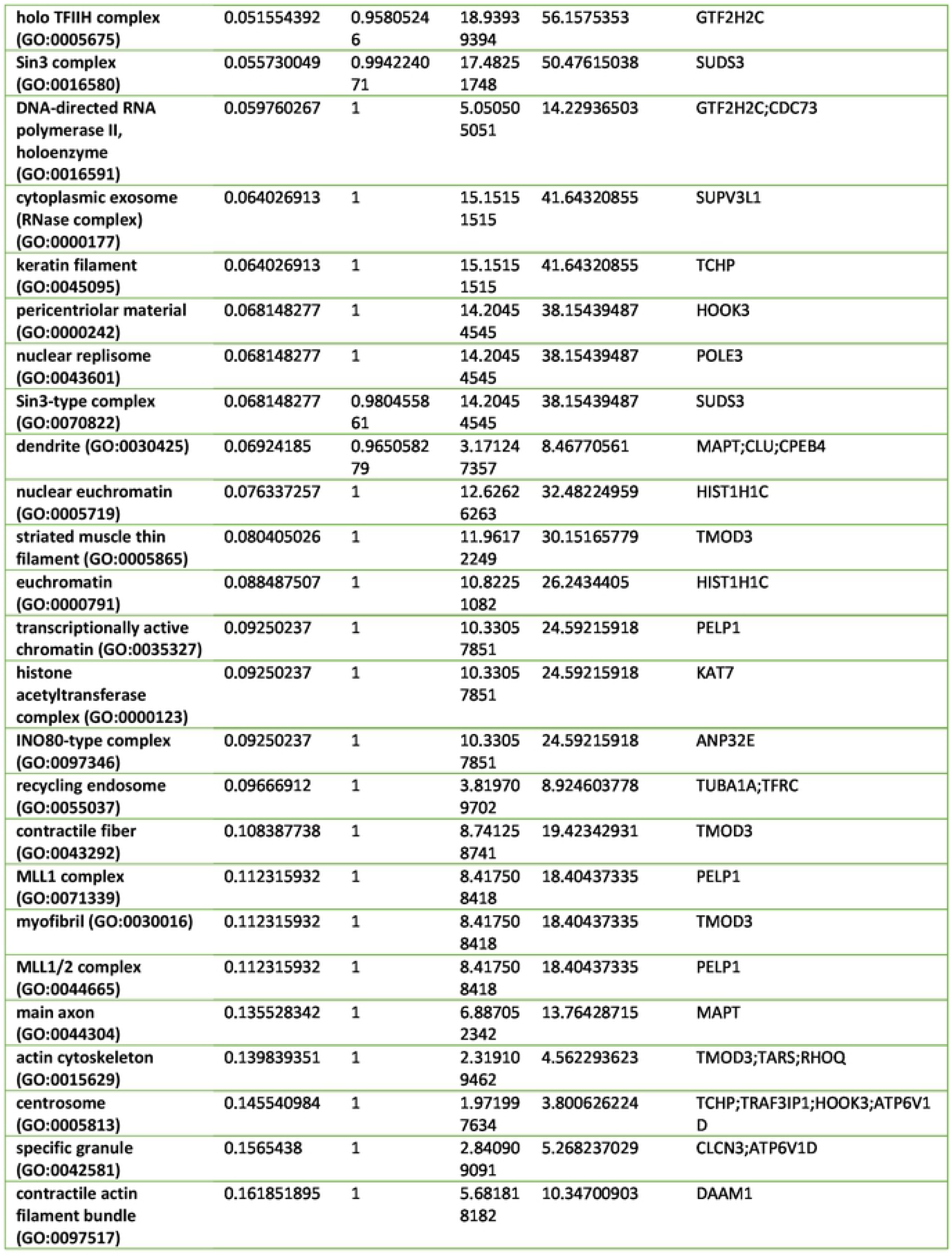
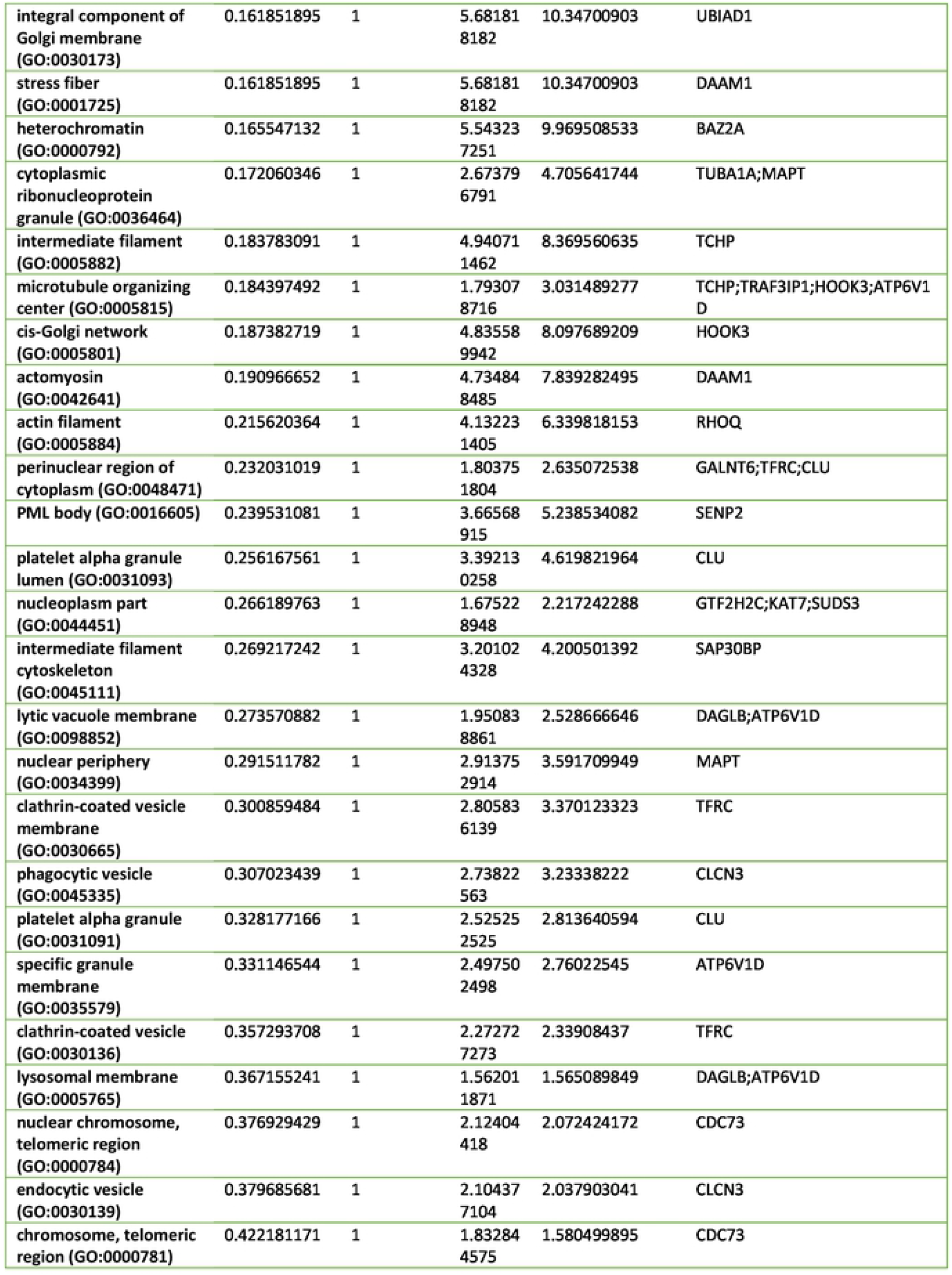
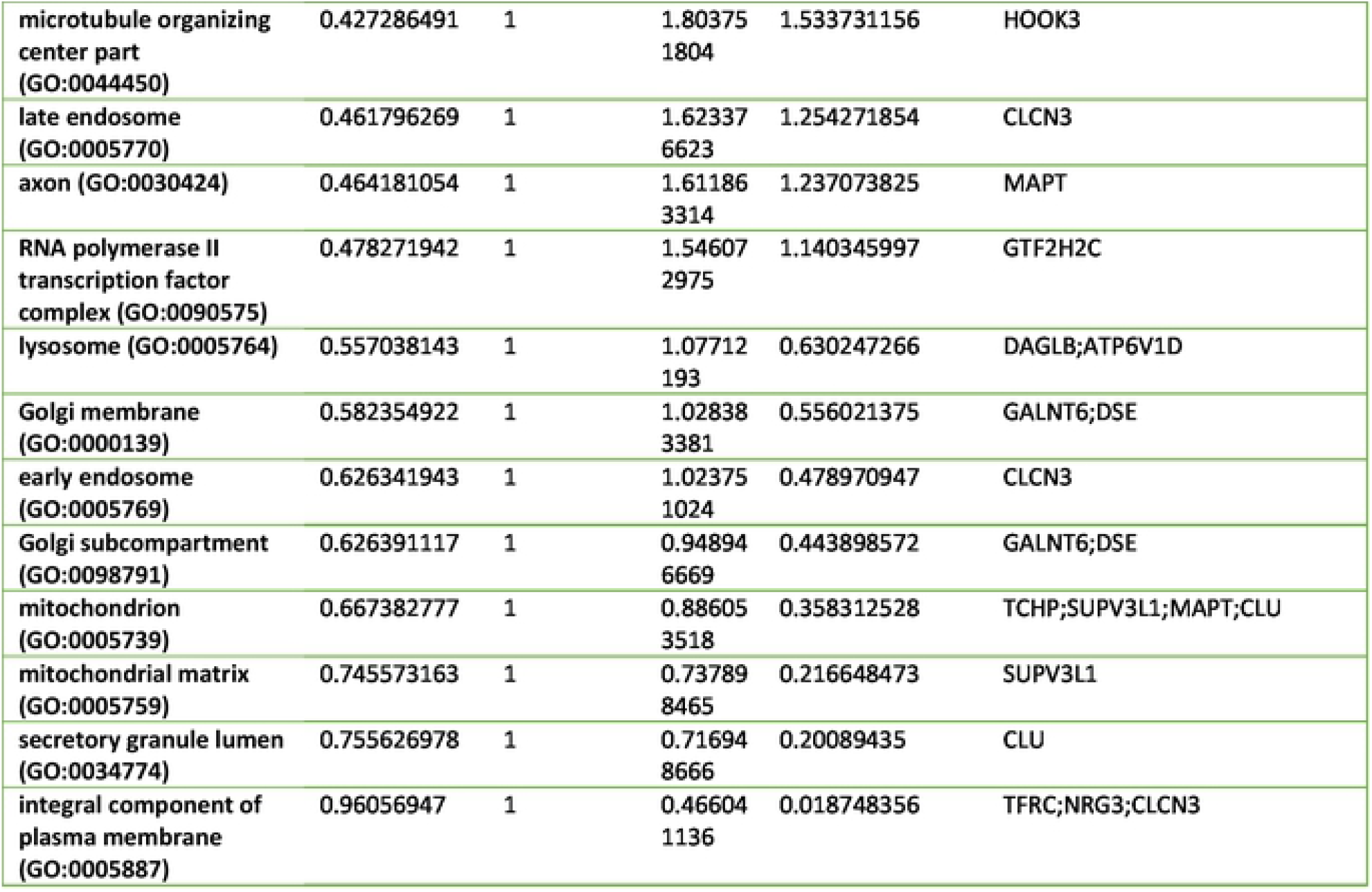
Up-Regulated Genes by WT Tau.

**Supplemental Table 2.** Down-Regulated Genes by WT Tau.

| **Term** | **P-value** | **Adjusted P-value** | **Odds Ratio** | **Combined Score** | **Genes** |
| --- | --- | --- | --- | --- | --- |
| axon initial segment (GO:0043194) | 0.013421829 | 1 | 74.07407407 | 319.3239135 | KCNQ2 |
| node of Ranvier (GO:0033268) | 0.017856968 | 1 | 55.55555556 | 223.6311935 | KCNQ2 |
| main axon (GO:0044304) | 0.048367756 | 1 | 20.2020202 | 61.19034096 | KCNQ2 |
| SCF ubiquitin ligase complex (GO:0019005) | 0.077961385 | 1 | 12.34567901 | 31.50051407 | KLHL11 |
| coated vesicle (GO:0030135) | 0.101264667 | 1 | 9.389671362 | 21.5025138 | STX6 |
| ribosome (GO:0005840) | 0.108009525 | 1 | 8.771929825 | 19.52224439 | NCK1 |
| perinuclear region of cytoplasm (GO:0048471) | 0.109697823 | 1 | 3.527336861 | 7.795505317 | STC2;STX6 |
| trans-Golgi network membrane (GO:0032588) | 0.120027161 | 1 | 7.843137255 | 16.62774289 | STX6 |
| clathrin-coated vesicle (GO:0030136) | 0.139709564 | 1 | 6.666666667 | 13.1212637 | STX6 |
| Golgi membrane (GO:0000139) | 0.141642689 | 1 | 3.016591252 | 5.895769726 | STX6;BLZF1 |
| Golgi subcompartment (GO:0098791) | 0.160952725 | 1 | 2.783576896 | 5.084605682 | STX6;BLZF1 |
| mitochondrion (GO:0005739) | 0.197703841 | 1 | 1.949317739 | 3.159815052 | TRUB1;OXCT1;PFDN2 |
| cullin-RING ubiquitin ligase complex (GO:0031461) | 0.238857152 | 1 | 3.683241252 | 5.273994836 | KLHL11 |
| trans-Golgi network (GO:0005802) | 0.243455603 | 1 | 3.603603604 | 5.091245694 | STX6 |
| nucleolus (GO:0005730) | 0.269438542 | 1 | 1.972386588 | 2.586617276 | UBE2T;UPF3A |
| early endosome (GO:0005769) | 0.284737465 | 1 | 3.003003003 | 3.772335434 | STX6 |
| endoplasmic reticulum lumen (GO:0005788) | 0.335058182 | 1 | 2.469135802 | 2.699879222 | STC2 |
| mitochondrial matrix (GO:0005759) | 0.37245128 | 1 | 2.164502165 | 2.13776849 | OXCT1 |
| integral component of plasma membrane (GO:0005887) | 0.377736622 | 1 | 1.367053999 | 1.330906484 | SLC26A2;KCNQ2;ICAM5 |

**Supplemental Table 3.** Up-Regulated Genes by P301L Tau.

| **Term** | **P-value** | **Adjusted P-value** | **Odds Ratio** | **Combined Score** | **Genes** |
| --- | --- | --- | --- | --- | --- |
| axolemma (GO:0030673) | 0.002996578 | 1 | 333.3333333 | 1936.761417 | MAPT |
| dendrite (GO:0030425) | 0.004890312 | 1 | 18.60465116 | 98.98603085 | NLGN1;MAPT |
| filopodium tip (GO:0032433) | 0.004989818 | 0.741819593 | 200 | 1060.071173 | NLGN1 |
| spanning component of membrane (GO:0089717) | 0.005487568 | 0.611863795 | 181.8181818 | 946.412758 | NLGN1 |
| nuclear speck (GO:0016607) | 0.009082553 | 0.810163748 | 13.51351351 | 63.53243152 | ITPKC;MAPT |
| microtubule cytoskeleton (GO:0015630) | 0.015239427 | 1 | 10.30927835 | 43.13267377 | FER;MAPT |
| main axon (GO:0044304) | 0.016381548 | 1 | 60.60606061 | 249.1878617 | MAPT |
| cytoskeleton (GO:0005856) | 0.026441482 | 1 | 7.692307692 | 27.94477859 | FER;MAPT |
| filopodium (GO:0030175) | 0.029604609 | 1 | 33.33333333 | 117.3275068 | NLGN1 |
| nuclear body (GO:0016604) | 0.036393121 | 1 | 6.472491909 | 21.44579618 | ITPKC;MAPT |
| nuclear periphery (GO:0034399) | 0.038330884 | 1 | 25.64102564 | 83.62818815 | MAPT |
| ribonucleoprotein granule (GO:0035770) | 0.039296097 | 1 | 25 | 80.91575164 | MAPT |
| axon (GO:0030424) | 0.068319537 | 1 | 14.18439716 | 38.0646738 | MAPT |
| RNA polymerase II transcription factor complex (GO:0090575) | 0.07113123 | 1 | 13.60544218 | 35.9622965 | FOS |
| cytoplasmic ribonucleoprotein granule (GO:0036464) | 0.081838782 | 1 | 11.76470588 | 29.44710636 | MAPT |
| microtubule (GO:0005874) | 0.100196316 | 1 | 9.523809524 | 21.91070345 | MAPT |
| polymeric cytoskeletal fiber (GO:0099513) | 0.105186383 | 1 | 9.049773756 | 20.38028436 | MAPT |
| nuclear chromatin (GO:0000790) | 0.119561652 | 1 | 7.90513834 | 16.78990615 | FER |
| actin cytoskeleton (GO:0015629) | 0.137676093 | 1 | 6.802721088 | 13.48878577 | FER |
| chromatin (GO:0000785) | 0.138551081 | 1 | 6.756756757 | 13.35483925 | FER |
| nuclear chromosome part (GO:0044454) | 0.179622583 | 1 | 5.102040816 | 8.759680555 | FER |
| nucleolus (GO:0005730) | 0.291015997 | 1 | 2.958579882 | 3.652003078 | TSPYL2 |
| mitochondrion (GO:0005739) | 0.409477333 | 1 | 1.949317739 | 1.740494599 | MAPT |
| integral component of plasma membrane (GO:0005887) | 0.532245452 | 1 | 1.367053999 | 0.862133316 | NLGN1 |

**Supplemental Figure 4.**
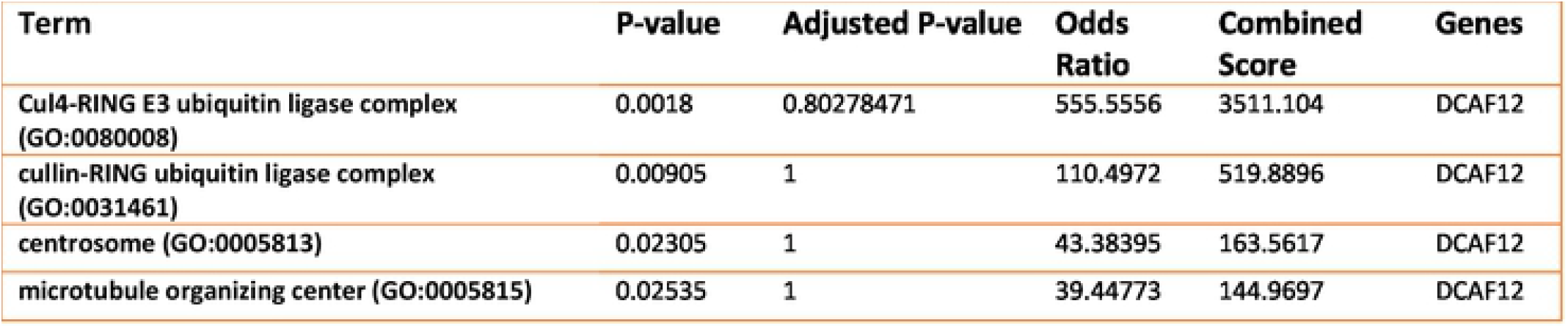
Down-Regulated Genes by P301L Tau.

## References

1. Sultan A, Nesslany F, Violet M, et al. Nuclear tau, a key player in neuronal DNA protection. The Journal of biological chemistry. 2011;286(6):4566–4575. doi:10.1074/jbc.M110.199976

2. Violet M, Delattre L, Tardivel M, et al. A major role for Tau in neuronal DNA and RNA protection in vivo under physiological and hyperthermic conditions. Frontiers in cellular neuroscience. 2014;8:84. doi:10.3389/fncel.2014.00084

3. Hua Q, He R. Tau could protect DNA double helix structure. Biochimica et biophysica acta. 2003;1645(2):205–211. doi:10.1016/s1570-9639(02)00538-1

4. Zhang X, Lin Y, Eschmann NA, et al. RNA stores tau reversibly in complex coacervates. PLoS biology. 2017;15(7):e2002183. doi:10.1371/journal.pbio.2002183

5. Kampers T, Friedhoff P, Biernat J, Mandelkow EM, Mandelkow E. RNA stimulates aggregation of microtubule-associated protein tau into Alzheimer-like paired helical filaments. FEBS letters. 1996;399(3):344–349. doi:10.1016/s0014-5793(96)01386-5

6. Monroy-Ramírez HC, Basurto-Islas G, Mena R, et al. Alterations in the nuclear architecture produced by the overexpression of tau protein in neuroblastoma cells. Journal of Alzheimer’s disease : JAD. 2013;36(3):503–520. doi:10.3233/JAD-122401

7. Montalbano M, McAllen S, Sengupta U, et al. Tau oligomers mediate aggregation of RNA-binding proteins Musashi1 and Musashi2 inducing Lamin alteration. Aging cell. Published online September 2019:e13035. doi:10.1111/acel.13035

8. Loomis PA, Howard TH, Castleberry RP, Binder LI. Identification of nuclear tau isoforms in human neuroblastoma cells. Proceedings of the National Academy of Sciences of the United States of America. 1990;87(21):8422–8426. doi:10.1073/pnas.87.21.8422

9. Shea TB, Cressman CM. A 26-30 kDa developmentally-regulated tau isoform localized within nuclei of mitotic human neuroblastoma cells. International journal of developmental neuroscience : the official journal of the International Society for Developmental Neuroscience. 1998;16(1):41–48. doi:10.1016/s0736-5748(97)00044-0

10. Ulrich G, Salvade A, Boersema P, et al. Phosphorylation of nuclear Tau is modulated by distinct cellular pathways. Scientific reports. 2018;8(1):17702. doi:10.1038/s41598-018-36374-4

11. Baquero J, Varriano S, Ordonez M, et al. Nuclear Tau, p53 and Pin1 Regulate PARN-Mediated Deadenylation and Gene Expression. Frontiers in Molecular Neuroscience. 2019;12:242. doi:10.3389/fnmol.2019.00242

12. Farmer KM, Ghag G, Puangmalai N, Montalbano M, Bhatt N, Kayed R. P53 aggregation, interactions with tau, and impaired DNA damage response in Alzheimer’s disease. Acta Neuropathologica Communications. 2020;8(1):132. doi:10.1186/s40478-020-01012-6

13. Montalbano M, McAllen S, Puangmalai N, et al. RNA-binding proteins Musashi and tau soluble aggregates initiate nuclear dysfunction. Nature Communications. 2020;11(1):4305. doi:10.1038/s41467-020-18022-6

14. Venkatramani A, Panda D. Regulation of neuronal microtubule dynamics by tau: Implications for tauopathies. International journal of biological macromolecules. 2019;133:473–483. doi:10.1016/j.ijbiomac.2019.04.120

15. Maina MB, Bailey LJ, Wagih S, et al. The involvement of tau in nucleolar transcription and the stress response. Acta neuropathologica communications. 2018;6(1):70. doi:10.1186/s40478-018-0565-6

16. Maina MB, Bailey LJ, Doherty AJ, Serpell LC. The Involvement of Abeta42 and Tau in Nucleolar and Protein Synthesis Machinery Dysfunction. Frontiers in cellular neuroscience. 2018;12:220. doi:10.3389/fncel.2018.00220

17. Lester E, Parker R. The Tau of Nuclear-Cytoplasmic Transport. Neuron. 2018;99(5):869–871. doi:10.1016/j.neuron.2018.08.026

18. Paonessa F, Evans LD, Solanki R, et al. Microtubules Deform the Nuclear Membrane and Disrupt Nucleocytoplasmic Transport in Tau-Mediated Frontotemporal Dementia. Cell reports. 2019;26(3):582-593.e5. doi:10.1016/j.celrep.2018.12.085

19. Tripathi T, Prakash J, Shav-Tal Y. Phospho-Tau Impairs Nuclear-Cytoplasmic Transport. ACS chemical neuroscience. 2019;10(1):36–38. doi:10.1021/acschemneuro.8b00632

20. Eftekharzadeh B, Daigle JG, Kapinos LE, et al. Tau Protein Disrupts Nucleocytoplasmic Transport in Alzheimer’s Disease. Neuron. 2018;99(5):925-940.e7. doi:10.1016/j.neuron.2018.07.039

21. Siano G, Varisco M, Caiazza MC, et al. Tau Modulates VGluT1 Expression. Journal of molecular biology. 2019;431(4):873–884. doi:10.1016/j.jmb.2019.01.023

22. Klein H-U, McCabe C, Gjoneska E, et al. Epigenome-wide study uncovers large-scale changes in histone acetylation driven by tau pathology in aging and Alzheimer’s human brains. Nature neuroscience. 2019;22(1):37–46. doi:10.1038/s41593-018-0291-1

23. Mansuroglu Z, Benhelli-Mokrani H, Marcato V, et al. Loss of Tau protein affects the structure, transcription and repair of neuronal pericentromeric heterochromatin. Scientific reports. 2016;6:33047. doi:10.1038/srep33047

24. Benhelli-Mokrani H, Mansuroglu Z, Chauderlier A, et al. Genome-wide identification of genic and intergenic neuronal DNA regions bound by Tau protein under physiological and stress conditions. Nucleic acids research. 2018;46(21):11405–11422. doi:10.1093/nar/gky929

25. Stewart M. Polyadenylation and nuclear export of mRNAs. The Journal of biological chemistry. 2019;294(9):2977–2987. doi:10.1074/jbc.REV118.005594

26. Gruber AJ, Zavolan M. Alternative cleavage and polyadenylation in health and disease. Nature Reviews Genetics. 2019;20(10):599–614. doi:10.1038/s41576-019-0145-z

27. Tian B, Hu J, Zhang H, Lutz CS. A large-scale analysis of mRNA polyadenylation of human and mouse genes. Nucleic Acids Research. 2005;33(1):201–212. doi:10.1093/nar/gki158

28. Shepard PJ, Choi E-A, Lu J, Flanagan LA, Hertel KJ, Shi Y. Complex and dynamic landscape of RNA polyadenylation revealed by PAS-Seq. RNA (New York, NY). 2011;17(4):761–772. doi:10.1261/rna.2581711

29. Rio DC, Ares MJ, Hannon GJ, Nilsen TW. Purification of RNA using TRIzol (TRI reagent). Cold Spring Harbor protocols. 2010;2010(6):pdb.prot5439. doi:10.1101/pdb.prot5439

30. Jaworski E, Routh A. ClickSeq: Replacing Fragmentation and Enzymatic Ligation with Click-Chemistry to Prevent Sequence Chimeras. Methods in molecular biology (Clifton, NJ). 2018;1712:71–85. doi:10.1007/978-1-4939-7514-3_6

31. Elrod ND, Jaworski EA, Ji P, Wagner EJ, Routh A. Development of Poly(A)-ClickSeq as a tool enabling simultaneous genome-wide poly(A)-site identification and differential expression analysis. Methods (San Diego, Calif). 2019;155:20–29. doi:10.1016/j.ymeth.2019.01.002

32. Routh A. DPAC: A Tool for Differential Poly(A)-Cluster Usage from Poly(A)-Targeted RNAseq Data. G3 (Bethesda, Md). 2019;9(6):1825–1830. doi:10.1534/g3.119.400273

33. Kim D, Paggi JM, Park C, Bennett C, Salzberg SL. Graph-based genome alignment and genotyping with HISAT2 and HISAT-genotype. Nature Biotechnology. 2019;37(8):907–915. doi:10.1038/s41587-019-0201-4

34. Kuleshov M v, Jones MR, Rouillard AD, et al. Enrichr: a comprehensive gene set enrichment analysis web server 2016 update. Nucleic Acids Research. 2016;44(W1):W90–W97. doi:10.1093/nar/gkw377

35. Miyasaka T, Shinzaki Y, Yoshimura S, et al. Imbalanced Expression of Tau and Tubulin Induces Neuronal Dysfunction in C. elegans Models of Tauopathy. Frontiers in Neuroscience. 2018;12:415. doi:10.3389/fnins.2018.00415

36. Bittermann E, Abdelhamed Z, Liegel RP, et al. Differential requirements of tubulin genes in mammalian forebrain development. PLOS Genetics. 2019;15(8):e1008243. https://doi.org/10.1371/journal.pgen.1008243

37. Kwon HS, Koh S-H. Neuroinflammation in neurodegenerative disorders: the roles of microglia and astrocytes. Translational Neurodegeneration. 2020;9(1):42. doi:10.1186/s40035-020-00221-2

38. Morales I, Jiménez JM, Mancilla M, Maccioni RB. Tau oligomers and fibrils induce activation of microglial cells. Journal of Alzheimer’s disease : JAD. 2013;37(4):849–856. doi:10.3233/JAD-131843

39. Karch CM, Goate AM. Alzheimer’s disease risk genes and mechanisms of disease pathogenesis. Biological psychiatry. 2015;77(1):43–51. doi:10.1016/j.biopsych.2014.05.006

40. Niday Z, Hawkins VE, Soh H, Mulkey DK, Tzingounis A v. Epilepsy-Associated KCNQ2 Channels Regulate Multiple Intrinsic Properties of Layer 2/3 Pyramidal Neurons. The Journal of neuroscience : the official journal of the Society for Neuroscience. 2017;37(3):576–586. doi:10.1523/JNEUROSCI.1425-16.2016

41. Short B, Preisinger C, Körner R, Kopajtich R, Byron O, Barr FA. A GRASP55-rab2 effector complex linking Golgi structure to membrane traffic. Journal of Cell Biology. 2001;155(6):877–884. doi:10.1083/jcb.200108079

42. Ito D, Walker JR, Thompson CS, et al. Characterization of stanniocalcin 2, a novel target of the mammalian unfolded protein response with cytoprotective properties. Molecular and cellular biology. 2004;24(21):9456–9469. doi:10.1128/MCB.24.21.9456-9469.2004

43. Bemben MA, Shipman SL, Nicoll RA, Roche KW. The cellular and molecular landscape of neuroligins. Trends in neurosciences. 2015;38(8):496–505. doi:10.1016/j.tins.2015.06.004

44. Brito-Moreira J, Lourenco M v, Oliveira MM, et al. Interaction of amyloid-β (Aβ) oligomers with neurexin 2α and neuroligin 1 mediates synapse damage and memory loss in mice. The Journal of biological chemistry. 2017;292(18):7327–7337. doi:10.1074/jbc.M116.761189

45. Dufort-Gervais J, Provost C, Charbonneau L, et al. Neuroligin-1 is altered in the hippocampus of Alzheimer’s disease patients and mouse models, and modulates the toxicity of amyloid-beta oligomers. Scientific reports. 2020;10(1):6956. doi:10.1038/s41598-020-63255-6

46. Craig AM, Kang Y. Neurexin-neuroligin signaling in synapse development. Current opinion in neurobiology. 2007;17(1):43–52. doi:10.1016/j.conb.2007.01.011

47. Helou J el, Bélanger-Nelson E, Freyburger M, et al. Neuroligin-1 links neuronal activity to sleep-wake regulation. Proceedings of the National Academy of Sciences of the United States of America. 2013;110(24):9974–9979. http://www.jstor.org/stable/42706115

48. Wu X, Morishita WK, Riley AM, Hale WD, Südhof TC, Malenka RC. Neuroligin-1 Signaling Controls LTP and NMDA Receptors by Distinct Molecular Pathways. Neuron. 2019;102(3):621-635.e3. doi:10.1016/j.neuron.2019.02.013

49. Lee S-H, Peng I-F, Ng YG, et al. Synapses are regulated by the cytoplasmic tyrosine kinase Fer in a pathway mediated by p120catenin, Fer, SHP-2, and beta-catenin. The Journal of cell biology. 2008;183(5):893–908. doi:10.1083/jcb.200807188

50. Patrón LA, Nagatomo K, Eves DT, et al. Cul4 ubiquitin ligase cofactor DCAF12 promotes neurotransmitter release and homeostatic plasticity. The Journal of cell biology. 2019;218(3):993–1010. doi:10.1083/jcb.201805099

51. Zheng C, Geetha T, Babu JR. Failure of ubiquitin proteasome system: risk for neurodegenerative diseases. Neuro-degenerative diseases. 2014;14(4):161–175. doi:10.1159/000367694

52. Paudel YN, Angelopoulou E, Piperi C, Othman I, Aamir K, Shaikh MF. Impact of HMGB1, RAGE, and TLR4 in Alzheimer’s Disease (AD): From Risk Factors to Therapeutic Targeting. Cells. 2020;9(2). doi:10.3390/cells9020383

53. Yatskevich S, Rhodes J, Nasmyth K. Organization of Chromosomal DNA by SMC Complexes. Annual review of genetics. 2019;53:445–482. doi:10.1146/annurev-genet-112618-043633

54. Aragon L, Martinez-Perez E, Merkenschlager M. Condensin, cohesin and the control of chromatin states. Current opinion in genetics & development. 2013;23(2):204–211. doi:10.1016/j.gde.2012.11.004

55. Geuens T, Bouhy D, Timmerman V. The hnRNP family: insights into their role in health and disease. Human genetics. 2016;135(8):851–867. doi:10.1007/s00439-016-1683-5

56. Lee Y-B, Chen H-J, Peres JN, et al. Hexanucleotide Repeats in ALS/FTD Form Length-Dependent RNA Foci, Sequester RNA Binding Proteins, and Are Neurotoxic. Cell Reports. 2013;5(5):1178–1186. doi:https://doi.org/10.1016/j.celrep.2013.10.049

57. Liu Y, Szaro BG. hnRNP K post-transcriptionally co-regulates multiple cytoskeletal genes needed for axonogenesis. Development. 2011;138(14):3079 LP – 3090. doi:10.1242/dev.066993

58. Lee EK, Kim HH, Kuwano Y, et al. hnRNP C promotes APP translation by competing with FMRP for APP mRNA recruitment to P bodies. Nature structural & molecular biology. 2010;17(6):732–739. doi:10.1038/nsmb.1815

59. Rajagopalan LE, Westmark CJ, Jarzembowski JA, Malter JS. hnRNP C increases amyloid precursor protein (APP) production by stabilizing APP mRNA. Nucleic acids research. 1998;26(14):3418–3423. doi:10.1093/nar/26.14.3418

60. Gami-Patel P, Bandopadhyay R, Brelstaff J, Revesz T, Lashley T. The presence of heterogeneous nuclear ribonucleoproteins in frontotemporal lobar degeneration with FUS-positive inclusions. Neurobiology of aging. 2016;46:192–203. doi:10.1016/j.neurobiolaging.2016.07.004

61. Mori K, Lammich S, Mackenzie IRA, et al. hnRNP A3 binds to GGGGCC repeats and is a constituent of p62-positive/TDP43-negative inclusions in the hippocampus of patients with C9orf72 mutations. Acta neuropathologica. 2013;125(3):413–423. doi:10.1007/s00401-013-1088-7

62. Low Y-H, Asi Y, Foti SC, Lashley T. Heterogeneous Nuclear Ribonucleoproteins: Implications in Neurological Diseases. Molecular Neurobiology. 2021;58(2):631–646. doi:10.1007/s12035-020-02137-4

63. Pelisch F, Pozzi B, Risso G, Muñoz MJ, Srebrow A. DNA damage-induced heterogeneous nuclear ribonucleoprotein K sumoylation regulates p53 transcriptional activation. The Journal of biological chemistry. 2012;287(36):30789–30799. doi:10.1074/jbc.M112.390120

64. Mandler MD, Ku L, Feng Y. A cytoplasmic quaking I isoform regulates the hnRNP F/H-dependent alternative splicing pathway in myelinating glia. Nucleic Acids Research. 2014;42(11):7319–7329. doi:10.1093/nar/gku353

65. Moujalled D, Grubman A, Acevedo K, et al. TDP-43 mutations causing amyotrophic lateral sclerosis are associated with altered expression of RNA-binding protein hnRNP K and affect the Nrf2 antioxidant pathway. Human Molecular Genetics. 2017;26(9):1732–1746. doi:10.1093/hmg/ddx093

66. Malik AM, Miguez RA, Li X, Ho Y-S, Feldman EL, Barmada SJ. Matrin 3-dependent neurotoxicity is modified by nucleic acid binding and nucleocytoplasmic localization. Taylor JP, ed. eLife. 2018;7:e35977. doi:10.7554/eLife.35977

67. Drummond E, Pires G, MacMurray C, et al. Phosphorylated tau interactome in the human Alzheimer’s disease brain. Brain : a journal of neurology. 2020;143(9):2803–2817. doi:10.1093/brain/awaa223

68. Campanella C, Pace A, Caruso Bavisotto C, et al. Heat Shock Proteins in Alzheimer’s Disease: Role and Targeting. International Journal of Molecular Sciences. 2018;19(9). doi:10.3390/ijms19092603

69. Sokpor G, Xie Y, Rosenbusch J, Tuoc T. Chromatin Remodeling BAF (SWI/SNF) Complexes in Neural Development and Disorders. Frontiers in molecular neuroscience. 2017;10:243. doi:10.3389/fnmol.2017.00243

70. Maeder CI, Kim J-I, Liang X, et al. The THO Complex Coordinates Transcripts for Synapse Development and Dopamine Neuron Survival. Cell. 2018;174(6):1436-1449.e20. doi:10.1016/j.cell.2018.07.046

71. Morris KJ, Corbett AH. The polyadenosine RNA-binding protein ZC3H14 interacts with the THO complex and coordinately regulates the processing of neuronal transcripts. Nucleic acids research. 2018;46(13):6561–6575. doi:10.1093/nar/gky446

72. Maziuk B, Ballance HI, Wolozin B. Dysregulation of RNA Binding Protein Aggregation in Neurodegenerative Disorders. Frontiers in molecular neuroscience. 2017;10:89. doi:10.3389/fnmol.2017.00089

73. Skliris A, Papadaki O, Kafasla P, et al. Neuroprotection requires the functions of the RNA-binding protein HuR. Cell death and differentiation. 2015;22(5):703–718. doi:10.1038/cdd.2014.158

74. Lu B, Gehrke S, Wu Z. RNA metabolism in the pathogenesis of Parkinson?s disease. Brain research. 2014;1584:105–115. doi:10.1016/j.brainres.2014.03.003

75. Bampton A, Gittings LM, Fratta P, Lashley T, Gatt A. The role of hnRNPs in frontotemporal dementia and amyotrophic lateral sclerosis. Acta Neuropathologica. 2020;140(5):599–623. doi:10.1007/s00401-020-02203-0

76. Couthouis J, Hart MP, Shorter J, et al. A yeast functional screen predicts new candidate ALS disease genes. Proceedings of the National Academy of Sciences of the United States of America. 2011;108(52):20881–20890. doi:10.1073/pnas.1109434108

77. Doyen C-M, An W, Angelov D, et al. Mechanism of polymerase II transcription repression by the histone variant macroH2A. Molecular and cellular biology. 2006;26(3):1156–1164. doi:10.1128/MCB.26.3.1156-1164.2006

78. Tian B, Manley JL. Alternative cleavage and polyadenylation: the long and short of it. Trends in biochemical sciences. 2013;38(6):312–320. doi:10.1016/j.tibs.2013.03.005

79. Koren SA, Hamm MJ, Meier SE, et al. Tau drives translational selectivity by interacting with ribosomal proteins. Acta Neuropathologica. 2019;137(4):571–583. doi:10.1007/s00401-019-01970-9

80. Younas N, Zafar S, Shafiq M, et al. SFPQ and Tau: critical factors contributing to rapid progression of Alzheimer’s disease. Acta neuropathologica. 2020;140(3):317–339. doi:10.1007/s00401-020-02178-y

81. Strang KH, Croft CL, Sorrentino ZA, Chakrabarty P, Golde TE, Giasson BI. Distinct differences in prion-like seeding and aggregation between Tau protein variants provide mechanistic insights into tauopathies. The Journal of biological chemistry. 2018;293(7):2408–2421. doi:10.1074/jbc.M117.815357

82. Aoyagi H, Hasegawa M, Tamaoka A. Fibrillogenic nuclei composed of P301L mutant tau induce elongation of P301L tau but not wild-type tau. The Journal of biological chemistry. 2007;282(28):20309–20318. doi:10.1074/jbc.M611876200

83. Barghorn S, Zheng-Fischhöfer Q, Ackmann M, et al. Structure, microtubule interactions, and paired helical filament aggregation by tau mutants of frontotemporal dementias. Biochemistry. 2000;39(38):11714–11721. doi:10.1021/bi000850r

